# Assessment of three-dimensional RNA structure prediction in CASP15

**DOI:** 10.1101/2023.04.25.538330

**Authors:** Rhiju Das, Rachael C. Kretsch, Adam J. Simpkin, Thomas Mulvaney, Phillip Pham, Ramya Rangan, Fan Bu, Ronan M. Keegan, Maya Topf, Daniel J. Rigden, Zhichao Miao, Eric Westhof

## Abstract

The prediction of RNA three-dimensional structures remains an unsolved problem. Here, we report assessments of RNA structure predictions in CASP15, the first CASP exercise that involved RNA structure modeling. Forty two predictor groups submitted models for at least one of twelve RNA-containing targets. These models were evaluated by the RNA-Puzzles organizers and, separately, by a CASP-recruited team using metrics (GDT, lDDT) and approaches (Z-score rankings) initially developed for assessment of proteins and generalized here for RNA assessment. The two assessments independently ranked the same predictor groups as first (AIchemy_RNA2), second (Chen), and third (RNAPolis and GeneSilico, tied); predictions from deep learning approaches were significantly worse than these top ranked groups, which did not use deep learning. Further analyses based on direct comparison of predicted models to cryogenic electron microscopy (cryo-EM) maps and X-ray diffraction data support these rankings. With the exception of two RNA-protein complexes, models submitted by CASP15 groups correctly predicted the global fold of the RNA targets. Comparisons of CASP15 submissions to designed RNA nanostructures as well as molecular replacement trials highlight the potential utility of current RNA modeling approaches for RNA nanotechnology and structural biology, respectively. Nevertheless, challenges remain in modeling fine details such as non-canonical pairs, in ranking among submitted models, and in prediction of multiple structures resolved by cryo-EM or crystallography.

## 1 Introduction

Soon after the establishment of the cloverleaf structure of transfer RNA,^1,2^ three-dimensional models of RNA structures appeared.^3,4^ However, it took more than ten years before the first refined experimental structures of the 76 nucleotide yeast tRNA^Phe^ were published.^5,6^ For many years, X-ray crystallographic structures of RNA nucleosides and nucleotides allowed us to grasp the fundamentals of RNA stereochemistry. After 1995, following progress in chemistry and X-ray technology, a steady stream of RNA structures with sizes equivalent to or larger than tRNAs, culminating with fully functional ribosome structures, revealed the many intricacies of RNA architectures. In parallel, computer programs for RNA modeling appeared (for overview, see ref.^7^). However, it was not until 2011 that a regular assessment of models, called RNA-Puzzles, was set up.^7,8^ The models for the RNA sequence of each RNA-Puzzle were collected prior to publications of the X-ray structures Since not enough targets were available for a short CASP-like season, the Puzzles were organized to occur right as the structures were solved (for those structures for which an agreement between the structural biologist and RNA-Puzzles organizers was made). Since then, several additional publications have reported the results of the RNA-Puzzles assessments.^9,10,11^ In 2021, it became clear that accelerations in RNA structure determination^12^ would allow enough targets for a single CASP season. Here we report on the first collaborative effort between CASP and RNA-Puzzles teams on a set of RNA targets. Following the success of AI-based tools in protein structure prediction^13^ and a surge of interest in RNA during the COVID pandemic,^14^ the hope of the organizers and assessors was to generate motivation and attention from protein modeling groups to develop and evaluate methods for RNA.

Between April and July of 2022, sequences of twelve RNA targets were received from experimental contributors and disseminated on the CASP website. Models were submitted by over 40 groups, and a double-blind assessment was carried out. Inspired by prior joint assessments by CAPRI and CASP for protein complexes (see, e.g. refs.^15–19)^, two assessments were carried out for RNA: one assessment was performed by the RNA-Puzzles team (Z. Miao & E. Westhof) and a completely independent analysis was performed by assessors nominated b the CASP organizers (R. Das and team). During a dedicated assessors’ meeting in October 2022, the two assessments’ results were critically compared, revealing a striking consensus in rankings and choice of top predictors, despite the use of distinct metrics and ranking schemes. Further analysis based on visual inspection of RNA-protein targets, direct comparison to cryogenic electron microscopy (cryo-EM) maps, and molecular replacement trials for targets solved by X-ray diffraction – catalyzed by the general CASP15 conference in December 2022 – revealed additional insights into the limitations and potential of current RNA 3D modeling, which are described here. The identification of accurate models also led to insights by CASP15 RNA experimental contributors and development of novel methods for cryo-EM model refinement, described in two separate papers co-submitted to the CASP15 special issue.^20,21^

## 2 Methods

### 2.1 Computation of RNA-Puzzles-style metrics

The RNA-puzzles-style assessment relied mainly on the Root Mean Square Deviation (RMSD) measure complemented by the Deformation Index (DI)^22^. The RMSD is the usual measure of distance between all atoms (excluding H atoms) of the two superimposed structures. The DI score complements the RMSD values by introducing features specific to RNA in the metric in the following way. The pairs formed by the nucleotides are identified, counted, and annotated in the experimental structure. They are broadly classified as either of the Watson-Crick complementary type (WC, comprising AU, GC, or GU pairs whose geometry are compatible with the standard Watson-Crick-Franklin double helix) or of the non-Watson-Crick type (NWC). The base–base network, i.e. WC, NWC, and stacking interactions in both reference and predicted models are extracted using the MC-Annotate^23^ tool. We then compute, for each of the three types of base-base interactions, the number of correctly predicted pairs, the true positive (TP), the number of predicted pairs with no correspondence in the reference model, the false positive (FP), and the number of pairs in the reference model that are not present in the predicted model, the false negative (FN). The Interaction Network Fidelity (INF) is then computed as the Matthews Correlation Coefficient, the geometric mean of the positive predictive value and sensitivity as in Gorodkin^24,25^:

The DI is then computed as: RMSD/INF. Several partial INF values (and respective DI) can be computed considering only the Watson-Crick (WC) base pairs (INF_WC_), the non-Watson-Crick (NWC) base pairs (INF_NWC_), both WC and NWC base pairs (INF_BPS_), or the stacking interactions (INF_STACK_). Finally, the Deformation Profile is a distance matrix computed as the average RMSD between the individual bases of the predicted and the reference models while superimposing each nucleotide of the predicted model over the corresponding nucleotide of the reference model one at a time. It is computed using the ‘‘dp.py’’ command from the ‘‘SIMINDEX’’ package^22^. For simplification, we also calculate the sum, mean and median of the deformation profile to account for the general accuracy of the prediction. The stereochemical correctness of the predicted models was evaluated with MolProbity^26^, which provides quality validation for 3D structures of proteins and nucleic acids. For the latter, MolProbity performs several automatic analyses, from checking the lengths of H-bonds present in the model to validating the compliance with the rotameric nature of the RNA backbone.^26,27^ As a single measure of stereochemical correctness, we chose the clash score, i.e., the number of all types of steric clashes per thousand residues.^28^ The assessment also considered the coordinate comparison metric TM-score as computed in RNA-Align^29^ and the Mean of Circular Quantities^30^ to assess accuracy in the torsion angle space. All the source codes and an example notebook are available at: https://github.com/RNA-Puzzles/RNA_assessment.

### 2.2 Computation of CASP-style metrics

Independently from the RNA-Puzzles-style computations, we assessed the accuracy of the submitted models in a manner closer to recent CASP assessments for protein structure prediction through Z_RNA_, a weighted Z-score average of several different assessment metrics. To perform the Z_RNA_ evaluation, we developed the casp-rna pipeline, which encompasses our workflow for data wrangling, job parallelization, and ranking visualizations. In consideration of RNA as a flexible molecule in which irregular loops may affect RMSD measures, Z_RNA_ explored additional metrics beyond RMSD to capture the global accuracy, local accuracy, and geometries of RNA. We selected the following tools for our ranking scheme: (1) US-align^31^, which was used to compute TM-score through a heuristic alignment approach improving on the original RNA-align^29^; (2) Local-Global Alignment^32^ which yielded GDT_TS, the average percentage of aligned C4’ atoms (rather than that of CL in proteins) at cutoffs of 1 Å, 2 Å, 4 Å, and 8 Å; (3) rna-tools^33^, a toolkit used to determine the accuracy of contact classifications among base stackings, Watson-Crick interactions, and non-canonical interactions. INF scores were calculated from interaction predictions dependent on ClaRNA^34^; (4) OpenStructure^35,36^, a framework used to find lDDT, a metric that measures structural similarity (unlike for proteins, our implementation of lDDT for RNA did not penalize for stereochemical violations); and (5) PHENIX, which reports a clashscore metric for all non-hydrogen bonded atom pairs that overlap worse than 0.4 Å^26,37^. For TM-score and GDT_TS, superposition of models and experimental models were calculated with default atoms for those packages, C3’ and C4’, respectively (repeating GDT_TS calculations with different atoms P, C3’, and C4’ gave negligible differences). Two alignment modes were considered for GDT_TS: a fixed residue-residue correspondence approach and an automated search for the best superposition, ignoring sequence; these gave nearly identical group rankings, so we opted for the former approach. INF scores were computed with ClaRNA to help increase robustness of base pair assignment for low resolution models; these values were slightly different than but highly correlated with INF scores computed with MC-Annotate, the tool normally used by RNA-Puzzles (**Supplemental Figure 1**).

Similar to the assessment of protein models in past CASP assessments, we employ a two-pass procedure for Z-scores^13,38^. For each target and for each of the considered metrics, the Z-score (difference with the mean, normalized by the standard deviation) was calculated by taking the mean and standard deviation for the best model from each group with respect to each considered metric. To prevent distortion from very poor outlier predictions, models with initial Z-scores that fall under a tolerance threshold of -2 were discarded, and the Z-scores were recomputed with the new mean and standard deviation. After this second pass, models with Z < -2 were re-assigned Z = -2. For Z-scores that involved linear combinations of multiple components (e.g., Z_RNA_), the Z-score values for individual components were then summed. To prevent penalization of novel methods that might give poor models for some targets, the sums of just the positive Z_RNA_ over all targets were used to make final rankings. For targets where experimentalists provided multiple conformations to either represent experimental uncertainty or bona fide conformational diversity (e.g., different copies in the crystallographic asymmetric unit or multiple conformations captured by cryo-EM^39^), predictor models were compared to all available experimental models. Groups were rewarded based on their best score. Code for the analysis of submitted models, assessment tools, and documentation using casp-rna are available as an open-source repository at https://github.com/DasLab/casp-rna. Metrics are also available for interactive viewing on the CASP15 website at https://predictioncenter.org/casp15/results.cgi?tr_type=rna.

### 2.3 Generation of simple template-based structures as comparison models

As baselines for the accuracy of predicted models, we prepared template-based structures generated using homology models with the rna_thread application in Rosetta 3 (version tag v2019.27-dev60818-134-g04678680f9c).^40^ For the CPEB3 ribozymes (R1107 and R1108), we generated template-based structures using the HDV ribozyme structure (PDB ID: 3NKB). We used residues 2-9, 11-39, 43-47, and 57-72 in this HDV ribozyme structure to model residues 3- 8, 10-43, and 54-69 in the CPEB3 ribozymes, avoiding loop residues that were not homologous between the structures. For the class I Pre-Q1 riboswitch, we compared the type III structure R1117 to template-based structures derived from the type I structure (PDB ID: 3Q50). We used residues 2-5, 7-20, and 23-33 in the type I Pre-Q1 riboswitch structure to model residues 2-30 in the target type III Pre-Q1 riboswitch structure, again avoiding loop residues that were not homologous between the models. Non-homologous residues were left out of these simple template-based structures.

### 2.4 Computation of map-to-model metrics for cryo-EM targets

All models for the 6 targets determined by cryo-EM (R1126, R1128, R1136, R1138, R1149, and R1156) were assessed directly against the experimental maps. The RNA-protein targets (R1189 and R1190) were excluded from this analysis because none of the predicted models for these targets fit sufficiently well into the density to give robust alignments, but in principle this analysis is compatible with RNA-protein targets. First, models were fit into maps using two approaches. Models were aligned to the reference models (built by experimentalists into density maps) using US-align^31^ and then fit locally using the command *fitmap* in ChimeraX^41^. We also tested an iterative phenix.dock_in_map^42^ procedure. For the well-fitting models, there was very little difference between these two methods and thus the *fitmap* method was selected. The following programs were used to measure the listed metrics, in all cases using default parameters (1) Phenix^42^, for cross-correlation of the map and model masked by the area around the model (CC_mask_), cross-correlation of the N highest density peaks in the model-generated map to the map (CC_volume_), cross-correlation of the N highest density peaks in the model-generated map and N highest density peaks in the map (CC_peaks_), and map-to-model Fourier shell correlation (FSC) values (N is the number of grid points inside the molecular mask); (2) TEMPy^43^, for cross-correlation coefficient (CCC), mutual information (MI), least-square fit (LSF), envelope score (ENV), and segment-based Mander’s overlap coefficient (SMOC); (3) ChimeraX^41^ and in-house script for atomic inclusion^44^ and density occupancy; and (4) MapQ^45^, for Q-score. An RMSD filter was selected for each target based on visual inspection. Ranking of all the models was carried out by Z-score, following the two-pass procedure described in section 2.2. Code for the analysis can be found at https://github.com/DasLab/CASP15_RNA_EM.

### 2.5 Scoring against X-ray data and molecular replacement (MR)

All models for the four targets determined by X-ray crystallography (R1107, R1108, R1116 and R1117) were assessed directly against the X-ray data by superimposing them on the target structure with RNAalign^29^ and calculating the Log Likelihood Gain (LLG) with respect to the diffraction data using Phaser^46^. For R1108 and R1117, with two RNA molecules in the asymmetric unit, the LLG was calculated for a single copy of the model ideally placed on chain A. A ranking of groups was derived from Z-scores computed from equal weighting of LLG, TFZ (translation-function Z-score from the model search), and CC (correlation coefficient of the map based on phases from the ideally placed model compared to the map computed by the experimentalists with their final phases). These ranking Z-scores were based on the same two-pass procedure as described in section 2.2.

Molecular Replacement was carried out using the CCP4 package^47^ via CCP4 Cloud^48^ and specifically the programs Phaser^46^ and MOLREP^49^. Map correlation coefficients were calculated with the phenix.get_cc_mtz_pdb tool^42^. MR strategies were chosen with reference to the accuracy achieved for different targets: highly accurate predictions typically succeed unmodified while extensive manual intervention can be required with poorer predictions. For R1117, the models were used unedited from all groups. For the other targets, where overall modeling was less accurate, different editing approaches were used with the models from group TS232 (AIchemy_RNA2). For R1107 and R1108, RNA model superposition was carried out with Theseus^50^ and nucleotides with higher structural variance values were removed in 10% intervals. The group 232 model_1 after removal of 10, 20, 30, 40 or 50% of nucleotides with highest structural divergence across the models was then used as a search model. MR also made use of models of the U1 small nuclear ribonucleoprotein A protein (U1ABD) component, which were generated using the AlphaFold 2^51^ network in its local ColabFold implementation^52^. For R1116, a version of Slice’N’Dice^53^ modified to work with RNA inputs was used to split model 1 from group TS232 into three structural segments using the Birch algorithm from the SciKit toolbox^54^.

## 3 Results

### 3.1 Classification of the difficulties and qualities of the targets

In **Table 1**, the twelve targets are gathered along with notes on protein and ligand binding, evidence for multiple conformations, and experimental technique and resolution. The difficulty was considered as “easy” when homologous structures were present in the PDB and as of “medium” difficulty when the structural similarity could be deduced due to similar functions (e.g., the CPEB3 ribozymes self-cleave like a ribozyme of known structure from hepatitis delta virus). Two targets were ranked as “difficult” since no homologous structures had been published and the number of nucleotides was larger than 120. Finally, a fourth “non-natural” category was considered for targets that were human-designed and not found in nature (and thus without homologous sequences), since it was not clear *a priori* whether these cases would be easy or difficult to model. The majority of targets (8) were solved by cryo-EM, with the rest (4) by X-ray crystallography.

**Table 1.** Summary and descriptions of the 12 RNA targets in CASP15.

| Name | Type | Length | Stoichiometry | Experimental method<sup>a</sup> (resol., Å) | Clash score, expt. | RNA type | Multiple conformations? | Difficulty | Best RMSD (Å) | Notable Groups |
| --- | --- | --- | --- | --- | --- | --- | --- | --- | --- | --- |
| R1107 | RNA<sup>b</sup> | 69 | A2 | X (2.8) | 5 | CPEB3 ribozyme (human) | (small differences with R1108) | Medium<sup>c</sup> | 4.5 | TS232 |
| R1108 | RNA<sup>b</sup> | 69 | A2 | X (2.2) | 2.4 | CPEB3 ribozyme (chimpanzee) | Two conformations in asymmetric unit | Medium<sup>c</sup> | 4.5 | TS232 |
| R1116 | RNA | 157 | A1 | X (3.0) | 20.1 | Cloverleaf RNA |  | Difficult | 4.8 | TS285 |
| R1117 | RNA / Ligand<sup>e</sup> | 30 | A1 | X (2.3) | 4.2 | PreQ<sub>1</sub> class I type III riboswitch |  | Easy<sup>d</sup> | 2.0 | TS287 |
| R1126 | RNA<sup>e</sup> | 363 | A1 | E (5.6) | 70.4 | Traptamer |  | Non natural | 8.9 | TS232 |
| R1128 | RNA | 238 | A1 | E (5.3) | 1.8 | Paranemic Crossover Triangle |  | Non natural | 4.3 | TS232 |
| R1136 | RNA<sup>e</sup> | 374 | A1 | E (4.5) | 0 | Apta-FRET | RNA with ligand bound and without. | Non natural | 7.2 | TS232 |
| R1138 | RNA | 720 | A1 | E (5.2) | 0.1<sup>f</sup> | 6HBC | Kinetically-trapped young and mature | Non natural | 7.8 | TS232 |
| R1149 | RNA | 124 | A1 | E (4.7) | 0.25 | SARS-CoV-2 SL5 | 10 models capturing modeling uncertainty. | Difficult | 6.9 | TS110 |
| R1156 | RNA | 135 | A1 | E (5.8) | 0.5 | BtCoV-HKU5 SL5 | 4 maps capturing flexibility; 10 models per map capturing modeling uncertainty. | Difficult | 5.4 | TS128 |
| R1189 | RNA/protein | 173 | A1B6 | E (3.8) | 2.6 | RsmZ-RsmA | Small differences in R1189/R1190 RNA | Difficult | 16.6 | TS229<sup>g</sup> |
| R1190 | RNA/protein | 173 | A1B4 | E (4.6) | 1.8 | RsmZ-RsmA | Small differences in R1189/R1190 RNA | Difficult | 16.0 | TS229<sup>g</sup> |

<sup>a</sup> X = X-ray crystallography, E = cryo-electron microscopy.
<sup>b</sup> Constructs for R1107 and R1108 both included engineered loops to complex with U1A protein, which aided crystallization; these were not noted as RNA/protein targets during the prediction season.
<sup>c</sup> Known to be related to HDV ribozyme (Salehi-Ashtiani, K., Lupták, A., Litovchick, A., & Szostak, J. W. (2006). A Genomewide Search for Ribozymes Reveals an HDV-Like Sequence in the Human CPEB3 Gene. *Science*, 313(5794), 1788–1792. https://doi.org/10.1126/science.1129308).
<sup>d</sup> Three types of PreQ<sub>1</sub> riboswitches are known in class I: high resolution X-ray structures of types I & II were known (PDB: 7REX (Schroeder, G. M., Cavender, C. E., Blau, M. E., Jenkins, J. L., Mathews, D. H., & Wedekind, J. E. (2022)) A small RNA that cooperatively senses two stacked metabolites in one pocket for gene control. *Nature Communications*, 13(1). https://doi.org/10.1038/s41467-021-27790-8), 3Q50 (Jenkins, J. L., Krucinska, J., McCarty, R. M., Bandarian, V. & Wedekind, J. E. Comparison of a preQ<sub>1</sub> riboswitch aptamer in metabolite-bound and free states with implications for gene regulation. *J. Biol. Chem.* 286, 24626–24637 (2011)).
<sup>e</sup> R1117, R1126 and R1136 were noted as RNA/ligand targets during prediction season. A K<sup>+</sup> ion in a G-quadruplex in R1126 and small molecules bound to aptamers displayed in R1136 were not well-resolved in their respective cryo-EM maps and not assessed. Assessment of the pre queuosine ligand in R1117 is included in the overall CASP15 assessment of ligand binding, described separately (Xavier Robin, Gabriel Studer, Janani Durairaj, Jerome Eberhardt, Torsten Schwede, and W. Patrick Walters, "Assessment of Protein-Ligand Complexes in CASP15", under revision).
<sup>f</sup> The model of the mature conformation has a clashscore of 0.09 and the top 10 CASP15 predictions matched this model better than the early conformation (clashscore 63.7).
<sup>g</sup> Similar RMSD predictions came from TS229, TS239 and TS439, all submitted by the same laboratory (Yang).

### 3.2 Assessment and ranking based on RNA-Puzzles metrics

The RNA-Puzzles assessment recognizes that RNA architecture results from a set of coherent interaction networks stabilizing a given fold. There are several interaction networks: the set formed by all Watson-Crick pairs, the set of contacts formed by the stacking between the bases, and finally the set formed by the non-Watson-Crick pairs, the interactions characteristic of tertiary folding. In a 3D structure, the set of Watson-Crick is not always the one predicted because in the folded structure, pairs at the extremities of the helical segments can either disappear or new ones can be formed. The correct choice of stacking between nucleotides or helices is critical for the overall global fold of the RNA. A wrong choice in the helices of the core can lead to very different folds from the native one. Finally, the appropriate positions and orientations of several elements allow for specific non-Watson-Crick pairs to form and lock in the native structure. An approximate association of helices may yield a molecular shape or envelope roughly similar to the native structure, but generally more open and much less compact than the native fold. In such cases, the key sequence conservations that maintain the actual native RNA fold are neither observed nor understood from the modeled structure. Therefore, in addition to using RMSD as a major metric for assessment, the analyses also included distinct metrics that are more sensitive to the interaction networks that comprise RNA.

**Table 1** gives the best RMSDs reached by the modeling groups for the 12 targets; they range between 2 Å and close to 17 Å, with many models being in the range between 4.3 Å and 8.3 Å. The trend follows the difficulty level of the targets. Interestingly, for the non-natural designed RNAs, the RMSDs reached are below 8.3 Å. It can be recalled that in a double stranded RNA helix, the average distance between two successive phosphate groups is 7 Å. However, broadly speaking, except for targets R1189 and R1190 (for which the RMSDs reached are beyond 16 Å, see **Table 1**), the overall folding shapes are reproduced, as can be seen in **Figure 1** where all targets are superimposed on the best predicted model as ranked by RMSD.

**FIGURE 1.**
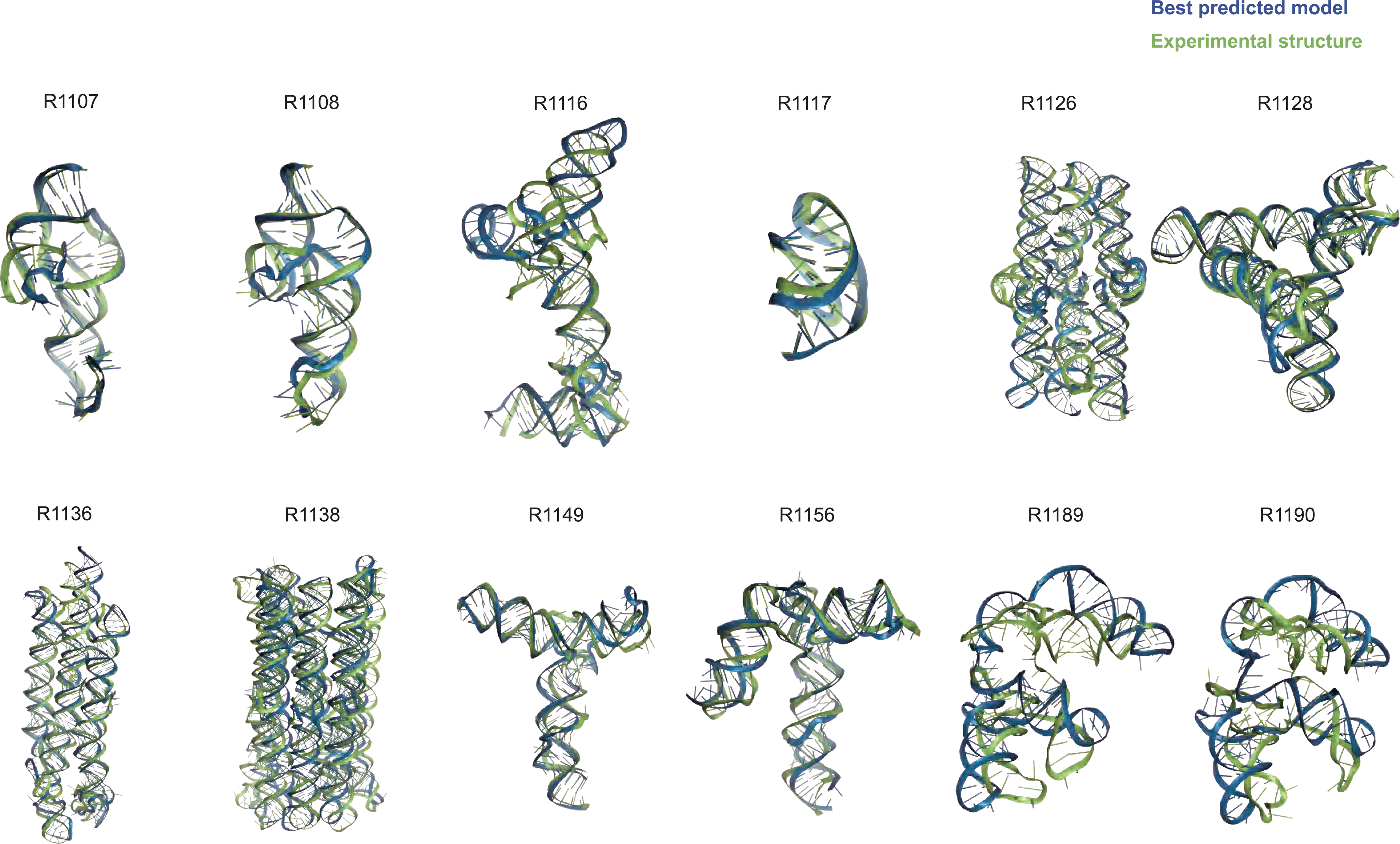
Overview of CASP15 RNA targets. Display of all CASP15 RNA targets (green) with the best-ranked model (blue) superimposed for each, chosen based on RMSD comparison of all five predicted models from all predictor groups compared to all available experimental structures. For ease of visualization of RNA global folds, protein binding and small molecule ligands (see **Table 1**) are not shown.

**Supplemental Table 1** presents the number of times that each of the modeling groups produced the 1st, 2nd, or 3rd best model as scored by the various metrics. Separate analyses are shown, based on the best of all five models from each predictor group and based solely on each groups’ model 1. Taking a weighted sum of these placements (with weights of 3, 2, and 1 assigned for placing 1st, 2nd, or 3rd) enables ranking of the groups. Whatever the way of counting or of scoring, even with methods that used metrics besides RMSD, two groups consistently reached the first and second ranks, TS232 (AIchemyRNA_2) and TS287 (Chen), respectively. The groups TS081 (RNApolis) and TS128 (GeneSilico) appear both at third positions. Considering those predictions with best RMSD that were ranked first amongst a set of all models submitted (up to five from each group), the groups TS232, TS287, TS081, and TS128 are the top four, with the other groups having weighted sums 50% lower. Among the latter, considering only at best RMSD rankings, TS229 (Yang-Server), TS416 (AIchemy_RNA), TS239 (Yang-Multimer), and TS439 (Yang) occupy the middle range.

### 3.3 Assessment based on CASP-style metrics

In a second assessment fully independent of the assessment based on RNA-Puzzles above, we explored the use of distinct metrics, largely drawn from assessment methods developed for proteins in previous CASP events and expanded here to RNA. For evaluating the global fold of predicted RNA structures, we computed the template modeling score (TM-score^29,31^) and the global distance test (GDT^32^). For the latter, we focused on the GDT score for tertiary structure (GDT_TS) rather than the high-accuracy GDT score (GDT_HA^55^) since the RNA models lacked nucleotide-level, much less atomic accuracy. To evaluate models’ local quality, to complement the RNA-specific INF score described in section 3.2, we used the Local Distance Difference Test (lDDT^35^) score, which compares distances between atoms that are nearby in the experimental structure to the distances between those atoms in the predicted structure and may generalize well between proteins and nucleic acids.

The global fold accuracy metrics (TM-score and GDT_TS) suggest that all targets, aside from the two RNA-protein complexes R1189 and R1190, elicited some predicted models that recovered correct global folds, based on criteria that have been previously discussed in the context of RNA template identification (TM-score > 0.45^29,31^, **Figure 2A**) or protein global fold assessment (GDT_TS > 45^56^, **Figure 2B**). We note that these criteria for ‘correct fold’ may not apply at the extremes of lengths for our RNA targets. On one hand, the “easy” PreQ_1_ riboswitch target (R1117) is small with only 30 nucleotides, and the TM-score values, which involve a length-dependent distance parameter, are much lower than GDT_TS values (**Figure 2A,B).** The accuracies reflected by GDT_TS match expected accuracies gauged by visual examination. On the other hand, models that visually captured correct folds for large designed RNA’s (R1126, R1128, R1136, R1138) were properly assigned high TM-scores, while GDT_TS scores were mostly lower than 45 (**Figure 2A-B**). For predictor models for a given target, the TM-score and

**FIGURE 2.**
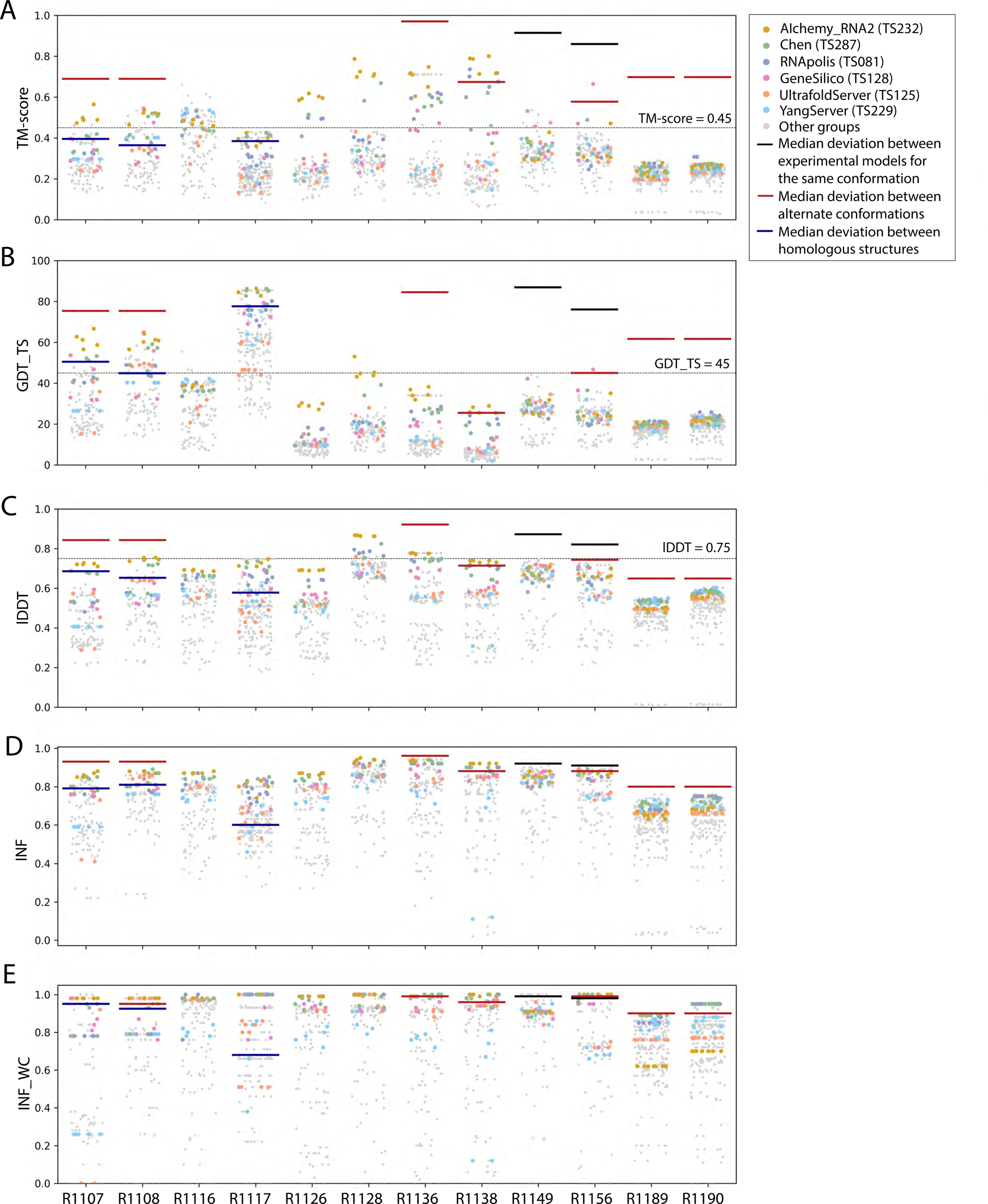
TM-score, GDT_TS, lDDT, INF, and INF_WC values for all targets. Scores for all models submitted for all targets are depicted (points are randomly jittered horizontally to aid visualization). Models from the four top performing groups and top two server groups are highlighted as colored points, and all other groups’ models are shown as gray points. Red lines indicate the median deviation between experimentally determined models for alternate conformations, black lines indicate the deviation between alternate models derived from experimental data for the same conformation, and blue lines indicate the deviation between homologous structures (see main text).

GDT_TS correlated well, but the relationship between the two varied across different targets (**Figure 3A**). The difference between GDT and TM-score is due to the distance cutoffs that the two metrics use. For example, TM-score applies a soft distance threshold *d*0 that depends on RNA length, which helps account for the flexibility of larger RNA’s.^29,31^ For R1138 (720 nt), *d*0 = 13.59 Å and most of the residues in a visually good model like R1138TS232_4 align within this threshold in the TM-score calculation. In contrast, GDT_TS uses fixed distance cutoffs of 1 Å, 2 Å, 4 Å, and 8 Å, and most of the RNA residues for the large molecules R1138TS232_4 do not align to the cryoEM structure within these thresholds (**Supplemental Figure 2)**. These comparisons suggest that TM-score and GDT are useful for ranking models for a given target but thresholds for ‘good’ TM-score and GDT may need recalibration for very small and very large RNA molecules, respectively.

**FIGURE 3:**
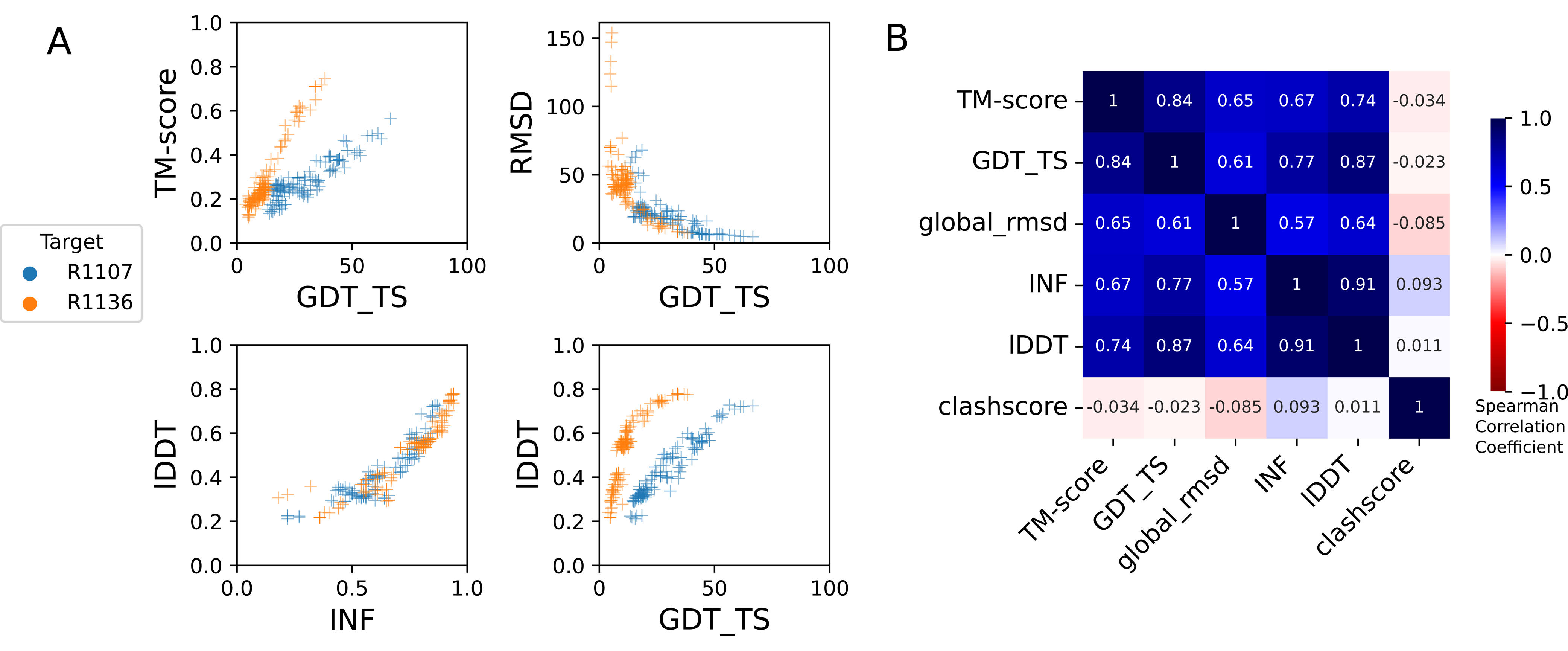
Comparison of assessment metrics for RNA targets. (A) Scores for all models for representative short target R1107 (blue) and long target R1136 (orange): top-left TM-score vs. GDT_TS, top-right RMSD vs. GDT_TS, to compare across global fold metrics; bottom-left lDDT vs. INF compares the two local metrics; and bottom-right lDDT vs. GDT_TS compares global fold to local metrics. **(B)** Average Spearman rank correlation coefficient (calculated separately per target, then averaged over all targets) between each pair of scores labeled on each row and column, colored by high correlation (dark blue), no correlation (white). RMSD and clashscore were multiplied by -1 before calculating the correlation so that higher scores correspond to better accuracy for all metrics.

As a metric for model quality that might generalize between protein and RNA, we considered lDDT. While not measuring global shape upon superposition, lDDT has been used as a primary accuracy indicator in numerous prediction contexts, including CAMEO, where a threshold of lDDT > 0.75 is used to denote a good match when comparing templates to target structures and to assign difficulty.^57,58^ Across all targets, lDDT values for best predictions ranged from 0.5 to 0.9, again with the lowest performance in RNA-protein complexes (**Figure 2C**). Interestingly, for the 10 RNA-only targets, CASP15 predictors achieved models with lDDT close to 0.75, and visually excellent models for the small, “easy” target R1117, the “medium” target R1108, and the “non-natural” and larger targets (R1128, R1136) achieved the 0.75 threshold. For future CASP, CAMEO, and other modeling challenges, lDDT may provide the most cleanly interpretable measure of accuracy, with a cutoff of 0.75 applicable across nucleic acids and proteins.

These CASP-inspired metrics correlated well with RNA-puzzle based metrics described in section 3.2. For global fold metrics, while RMSD and GDT_TS are not linearly correlated (**Figure 3A**), they have positive rank-based correlation (Spearman correlation coefficient 0.61, **Figure 3B**). The local interaction metrics, INF and lDDT, correlate excellently (Spearman correlation coefficient 0.91, **Figure 3B**) in what seems to be a near-linear and size-independent relationship (**Figure 3A**). This is a remarkably strong correlation; INF focuses on a selection of RNA-specific interactions while lDDT compares all heavy-atom distances for atom pairs that are within 15 Å in the experimental structure, a similar length scale to the distances across base pairs monitored by INF. This observation suggests that lDDT may capture the subset of interactions measured in INF while allowing generalization across protein and nucleic acids. Finally, if we compare global fold accuracy metrics with more local accuracy metrics, we still maintain a positive correlation (Spearman correlation coefficient 0.67-0.87, **Figure 3B**), however the relationship is non-linear; the more local metrics like lDDT are able to discriminate models with low accuracy while global fold metrics like GDT_TS are better able to discriminate the high accuracy models (**Figure 3A**).

To provide a more quantitative threshold for good model accuracy for each target, we sought to estimate the deviation between experimentally determined structures. Where possible, we measured the deviation in TM-score, GDT_TS, INF, INF_WC, and lDDT between distinct experimentally captured conformations (red lines in **Figure 2**). More specifically, we compared the following structure pairs in targets with multiple conformations (see also **Table 1**): the point-mutations for the CPEB3 ribozyme^59^ (R1107 vs R1108), the apo and holo structures of the aptamer Apta-FRET^60^ (R1136), the young and mature conformations of 6HBC^61^ (R1138), the four cryo-EM classes for the SL5 domain of the bat coronavirus HKU5 (R1156), and finally the RNA structures for the RsmZ-RsmA RNA-protein complexes with six vs. four proteins bound (R1189 vs R1190). In addition, for two cases, we measured the deviation between different models derived from the same experimental data (black lines in **Figure 2**), comparing distinct models built into the same cryo-EM density maps for the SL5 domains of SARS-CoV-2 (R1149) and BtCov-HKU5 (R1156). Finally, in three cases, we measured the deviation between homologous models, comparing residues that are homologous between previously solved structures and the target molecule (blue lines in Figure 2): the CPEB3 ribozyme versus the HDV ribozyme^62^ (R1107 and R1108 vs PDB ID 3NKB), and the class I type III Pre-Q1 riboswitch versus the class I type I Pre-Q1 riboswitch^63^ (R1117 vs PDB ID 3Q50). In all cases with available homologous structures (R1107, R1108, and R1117), predicted models surpassed TM-score, GDT_TS, lDDT, INF, and INF_WC values of models derived by directly using homologous structures. In some cases (R1138, R1156), predicted models reached TM-score, GDT_TS, and lDDT values comparable to the deviation between distinct experimentally determined conformations (red lines, **Figure 2**), though in no case were there models whose accuracies exceeded the experimental precision expected for a single captured conformation (black lines, **Figure 2**).

To rank the performance of predictors, we developed a Z-score metric that enabled combined evaluation of models’ global fold, local accuracy, and stereochemical correctness. Our global fold accuracy scores included the TM-score and GDT_TS, our more local accuracy scores consisted of INF and lDDT, and our stereochemical correctness scores were based on clashscore^28^, which has been used widely for both protein and RNA structural assessment. We used the following weighted sum of scores:

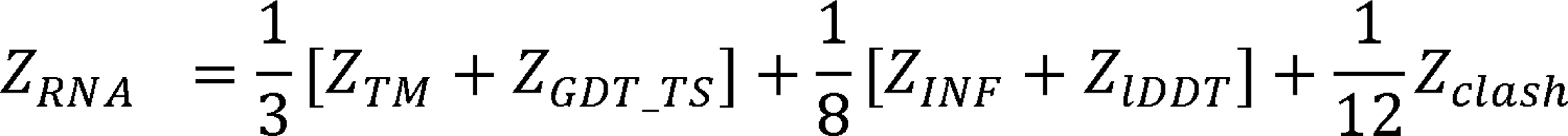

Because we did not expect atomically accurate models in this first RNA round of CASP, we chose to reward models that recover the global fold (high weight for TM-score and GDT_TS terms) compared to those that recover local details (low weight for local environment scores) or produce correct nucleotide geometries (low weight for clashscore). Each group’s Z-score for a given target was computed using their best predicted model, and groups’ total scores were calculated as the sum of all positive Z-scores across all targets (**Figure 4A**). The top performing predictor groups based on this combined Z-score ranking were AIchemy_RNA2 (TS232), Chen (TS287), RNAPolis (TS081), and Genesilico (TS128). These were the same groups as the top four highlighted by the independent analysis by the RNA-puzzles-style assessment.

**FIGURE 4.**
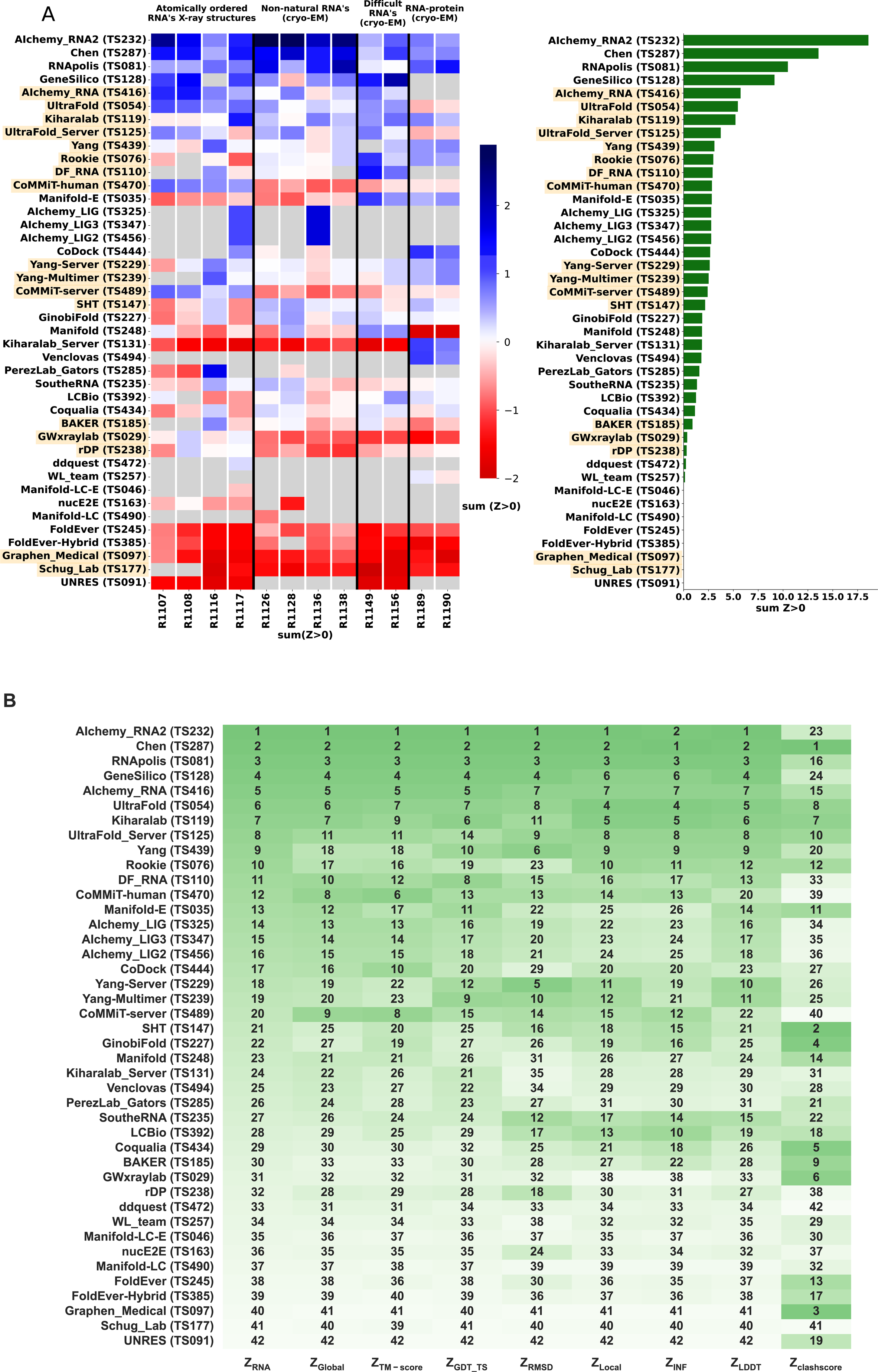
CASP-style Z-score based rankings. (A) Heatmap of groups ranked by Z_RNA_. Groups that used deep learning, as reported in the participant’s abstract to CASP15, are indicated in orange. The summation of positive two-pass Z-scores for each of the 12 targets is summarized in the barplot (right). Groups are ordered by their Z_RNA_ rankings. **(B)** Robustness of ranking to different choices in assessment. Columns show group rankings based on subsets of the Z_RNA_ score or individual metrics; coloring reflects rankings under each metric.

Interestingly, the top four groups did not include any server submissions; the top-ranked servers (Ultrafold-server, TS125; and Yang-server, TS229) placed at positions 8 and 9, and gave Z-scores that were more than three-fold lower than the top two predictor groups. We note that these top server submissions additionally exhibited secondary structures (Watson-Crick base-pairing) with lower accuracy than some other top predictors, as measured by INF_WC (orange and cyan points, Figure 2), suggesting that there is room for improvement in automated prediction of secondary structure. Furthermore, based on abstracts collected for the CASP15 conference, while the majority of CASP15 RNA predictors groups tested deep learning methods (orange highlights in **Figure 4A**), the top 4 RNA groups did not use deep learning approaches (see also articles by RNA predictor groups co-submitted for the CASP15 special issue^64^; and https://predictioncenter.org/casp15/doc/presentations/Day3/).

To better understand uncertainties in the rankings, we repeated the Z-score analysis using sub- components of the Z-score. Ranking groups by the two “global fold” terms (GDT_TS and TM- score) alone or in combination, or using RMSD, gave rankings with the same top four groups, up to some switching of third and fourth place (**Figure 4B** and **Table 2**). Use of the more local accuracy terms (lDDT and INF) retained the same top three predictor groups, with some groups switching in ranks of the groups after the top three. After the top four, the rankings are less consistent, which is not surprising given the small numerical score differences in these placements (**Figure 4A** and **Table 2**). Ranking groups by clashscore alone did not correlate with the other rankings (**Figure 3B** and **Figure 4B**), presumably because different predictors used somewhat different refinement schemes and were not told *a priori* that they would be assessed on clashscore. Overall, the ranking of the top four groups in CASP RNA structure modeling was robust to changes in metrics used and across two independent assessments.

**Table 2.** Z-scores for predictor groups using different combinations of assessment metrics.

| **Groups** | **Z<sub>RNA</sub>** | **Z<sub>TM-score</sub>** | **Z<sub>GDT_TS</sub>** | **Z<sub>RMSD</sub>** | **Z<sub>INF</sub>** | **Z<sub>IDDT</sub>** | **Z<sub>clashscore</sub>** |
| --- | --- | --- | --- | --- | --- | --- | --- |
| **Alchemy_RNA2 (TS232)** | 18.58 | 22.17 | 22.89 | 15.17 | 12.07 | 15.12 | 2.13 |
| **Chen (TS287)** | 13.56 | 15.49 | 13.70 | 13.59 | 12.46 | 13.73 | 6.74 |
| **RNapolis (TS081)** | 10.48 | 11.83 | 11.69 | 10.41 | 9.80 | 10.26 | 3.84 |
| **GeneSilico (TS128)** | 9.14 | 11.24 | 10.64 | 8.45 | 7.53 | 8.74 | 2.06 |
| **Alchemy_RNA (TS416)** | 5.73 | 6.25 | 5.72 | 6.36 | 6.55 | 6.21 | 4.07 |
| **UltraFold (TS054)** | 5.45 | 4.62 | 5.52 | 6.33 | 9.34 | 8.58 | 5.46 |
| **Kiharalab (TS119)** | 5.21 | 3.83 | 5.61 | 5.64 | 9.29 | 7.66 | 6.11 |
| **UltraFold_Server (TS125)** | 3.73 | 3.44 | 3.41 | 6.04 | 6.07 | 5.38 | 5.21 |
| **CoMMiT-human (TS470)** | 3.09 | 2.50 | 3.78 | 6.50 | 5.36 | 5.31 | 3.12 |
| **DF_RNA (TS110)** | 2.98 | 2.80 | 3.19 | 1.99 | 4.32 | 4.18 | 4.81 |
| **Yang-Server (TS229)** | 2.91 | 3.30 | 4.31 | 3.01 | 3.41 | 3.52 | 0.42 |
| **CoMMiT-server (TS489)** | 2.85 | 5.36 | 3.52 | 3.75 | 4.28 | 2.88 | 0.00 |
| **Alchemy_LIG (TS325)** | 2.83 | 2.73 | 3.76 | 2.12 | 2.13 | 3.09 | 5.04 |
| **Alchemy_LIG3 (TS347)** | 2.77 | 3.21 | 3.26 | 2.35 | 2.30 | 2.93 | 0.16 |
| **Alchemy_LIG2 (TS456)** | 2.77 | 3.21 | 3.26 | 2.35 | 2.30 | 2.93 | 0.16 |
| **Yang-Multimer (TS239)** | 2.77 | 3.21 | 3.26 | 2.35 | 2.30 | 2.93 | 0.16 |
| **Yang (TS439)** | 2.67 | 3.53 | 2.80 | 1.04 | 3.07 | 2.57 | 1.52 |
| **Manifold (TS248)** | 2.62 | 2.19 | 3.65 | 6.92 | 3.15 | 5.01 | 1.54 |
| **GinobiFold (TS227)** | 2.51 | 2.05 | 3.98 | 5.82 | 3.05 | 4.62 | 1.68 |
| **SHT (TS147)** | 2.41 | 4.34 | 3.29 | 3.52 | 4.28 | 2.76 | 0.00 |
| **BAKER (TS185)** | 2.16 | 2.38 | 1.98 | 2.68 | 3.69 | 2.77 | 6.68 |
| **Rookie (TS076)** | 1.86 | 2.40 | 1.43 | 1.61 | 3.45 | 2.37 | 6.66 |
| **SoutheRNA (TS235)** | 1.85 | 2.24 | 1.93 | 0.82 | 2.09 | 2.50 | 4.15 |
| **Manifold-E (TS035)** | 1.84 | 1.81 | 2.24 | 0.36 | 2.00 | 1.56 | 0.60 |
| **Coqualia (TS434)** | 1.78 | 1.79 | 2.21 | 0.37 | 1.60 | 1.49 | 0.73 |
| **PerezLab_Gators (TS285)** | 1.58 | 1.78 | 2.06 | 1.55 | 1.07 | 0.98 | 2.35 |
| **LCBio (TS392)** | 1.34 | 1.94 | 1.98 | 4.10 | 3.82 | 2.99 | 2.19 |
| **rDP (TS238)** | 1.25 | 1.81 | 1.21 | 2.67 | 4.52 | 2.90 | 3.54 |
| **GWxraylab (TS029)** | 1.14 | 1.53 | 0.63 | 1.78 | 3.18 | 2.16 | 6.64 |
| **CoDock (TS444)** | 0.89 | 0.55 | 1.10 | 1.31 | 2.66 | 1.60 | 5.26 |
| **ddquest (TS472)** | 0.36 | 1.10 | 0.78 | 0.42 | 0.00 | 0.13 | 6.11 |
| **nucE2E (TS163)** | 0.30 | 1.57 | 1.31 | 2.67 | 0.70 | 2.09 | 0.00 |
| **Manifold-LC-E (TS046)** | 0.23 | 1.33 | 0.12 | 0.40 | 0.00 | 0.02 | 0.00 |
| **Venclovas (TS494)** | 0.13 | 0.27 | 0.13 | 0.00 | 0.52 | 0.00 | 0.73 |
| **WL_team (TS257)** | 0.00 | 0.00 | 0.00 | 0.28 | 0.00 | 0.00 | 0.71 |
| **FoldEver (TS245)** | 0.00 | 0.21 | 0.05 | 1.95 | 0.00 | 0.36 | 0.00 |
| **FoldEver-Hybrid (TS385)** | 0.00 | 0.00 | 0.00 | 0.00 | 0.00 | 0.00 | 0.48 |
| **Kiharalab_Server (TS131)** | 0.00 | 0.05 | 0.00 | 0.98 | 0.00 | 0.00 | 4.17 |
| **Graphen_Medical (TS097)** | 0.00 | 0.00 | 0.00 | 0.29 | 0.00 | 0.00 | 3.78 |
| **UNRES (TS091)** | 0.00 | 0.00 | 0.00 | 0.00 | 0.00 | 0.00 | 6.68 |
| **Manifold-LC (TS490)** | 0.00 | 0.00 | 0.00 | 0.00 | 0.00 | 0.00 | 0.00 |
| **Schug_Lab (TS177)** | 0.00 | 0.00 | 0.00 | 0.00 | 0.00 | 0.00 | 3.54 |

### 3.4 Detailed assessment for RNA-protein complexes

The poorer predictions and the presence of RNA-protein contacts for the two RNA-protein complexes RT1189 and RT1190 largely precluded useful accuracy rankings from the metrics described above, so we carried out a detailed visual assessment for these targets. This assessment involved checking whether predictions had the right nucleotide – amino acid contacts and then visually assessing whether the fold was correct. For the contact-based analysis, a contact was defined as any pair of nucleotide and amino acid containing atoms within 5 Å of one another. The Matthews Correlation Coefficient (MCC) was used to score the contacts made by the predictions against those of the targets. The distribution of scores is shown in **Figure 5A**. The highest scoring model from each group with MCC scores above 0.1 (roughly the beginning of the non-zero peak in the distribution) were then visually assessed.

**FIGURE 5.**
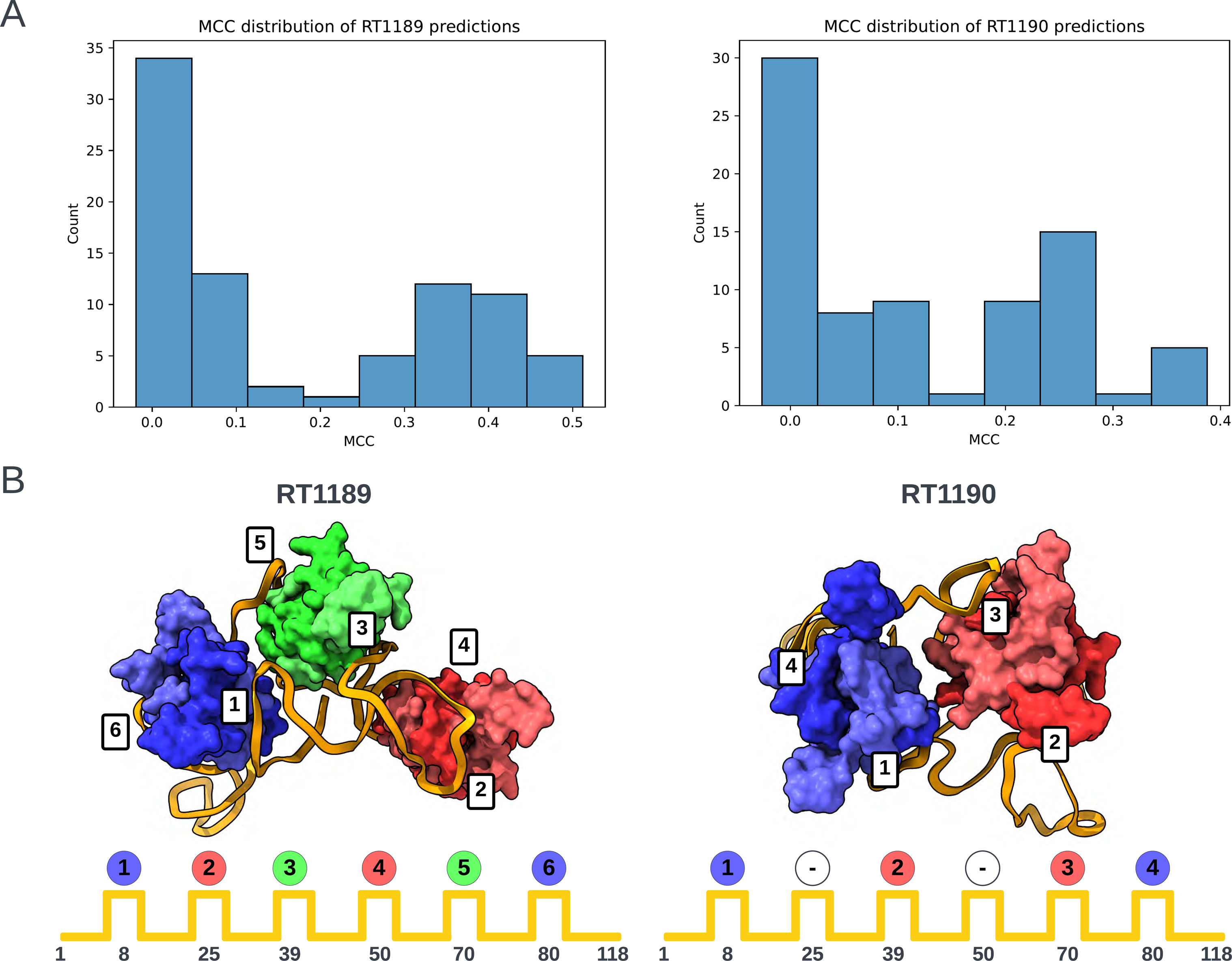
Folding pattern analysis of RNA-protein complexes. (A) Histograms of Matthews Correlation Coefficients (MCC) for RNA-protein contact accuracy in the two RNA-protein targets RT1189 and RT1190 (RsmZ-RsmA RNA-protein complexes). **(B)** Scheme for classifying the folding pattern of RNA based on order of protein contacts to RNA. Each dimer is assigned a color based on the order it was visited in. Experimental cryo-EM structures are shown at top with positions of binding on RNA diagrammed below.

For the RNA folding pattern analysis, we needed to establish a well-defined descriptor for the RNA-protein binding arrangement that was not dependent on superposition (which was difficult for all the models). This was achieved by coloring each protein by the regions of interaction in the RNA with the lowest order. Region order was determined by RNA sequence position (where 5’ is low). Using this scheme, the colors blue (B), then red (R), then green (G) were assigned to the three RsmA homodimers in RT1189 – and this pattern was compared for each model against the experimental structure (folding pattern: BRGRGB). In the case of RT1190, which involved only two RsmA homodimers, not all six regions of the RNA were bound; in particular, the regions of the RNA at approximately nucleotides 25 and 50 should not interact with a dimer.

For RT1189, no models exhibited the correct folding pattern for interacting with the 6 RsmA proteins (**Table 3**). For RT1190 (folding pattern string: B-R-RB), the best model according to the MCC score (MCC = 0.39) predicted the non-interacting RNA regions correctly (‘-’ in **Table 3**) but the RNA-protein contacts were made in the wrong order (B-R-BR). Many of the lower scoring models (MCC = 0.21-0.29), did contain interacting regions in the correct order but misplaced the non-interacting regions. As judged by this MCC contact-based score supplemented by protein-binding folding pattern analysis, TS119 (Kiharalab) and TS329 (LCBio) produced top-3 models for both targets (**Table 3**). In contrast, ranking based purely on RNA RMSD highlighted models from TS229 and other models from the Yang laboratory (**Table 1**); these models were less satisfactory from the point of view of protein-RNA contacts, showing the importance of complementary analyses in ranking these very difficult targets.

**Table 3.** Matthews correlation coefficients and folding pattern of the best model from each group with an MCC greater than 0.1. The symbols B, R, G and ‘-’ indicate blue, red, green and unbound regions as per Figure 5B.

| Target | Group ID | Group Name | Model | Contact MCC | Folding pattern |
| --- | --- | --- | --- | --- | --- |
| **RT1189**<br>**(native folding pattern: BRGRGB)** | 119 | Kiharalab | 3 | 0.51 | BRGRBG |
|  | 232 | Alchemy_RNA2 | 2 | 0.41 | BRGRBG |
|  | 392 | LCBio | 2 | 0.41 | BRGRBG |
|  | 444 | CoDock | 3 | 0.39 | BRGRBG |
|  | 439 | Yang | 2 | 0.38 | BRGRBG |
|  | 131 | Kiharalab_Server | 2 | 0.37 | BRGRBG |
|  | 494 | Venclovas | 4 | 0.32 | BRGRBG |
|  | 434 | Coqualia | 1 | 0.26 | 1 hexamer |
|  | 185 | BAKER | 1 | 0.18 | BRGB |
| **RT1190**<br>**(native folding pattern: B-R-RB)** | 392 | LCBio | 5 | 0.39 | B-R-BR |
|  | 119 | Kiharalab | 2 | 0.29 | BR-RB- |
|  | 444 | CoDock | 5 | 0.27 | BR-RB- |
|  | 494 | Venclovas | 4 | 0.26 | BR-RB- |
|  | 131 | Kiharalab_Server | 5 | 0.26 | BR-RB- |
|  | 232 | Alchemy_RNA2 | 4 | 0.21 | BR-RB- |
|  | 434 | Coqualia | 1 | 0.17 | Dimers conjoined |
|  | 035 | Manifold-E | 1 | 0.12 | Protein separated from RNA |
|  | 227 | GinobiFold | 1 | 0.10 | Dimers conjoined |

### 3.5 Ranking based on direct comparison to cryo-EM maps

The ‘native’ experimental models built from RNA cryo-EM maps may be particularly susceptible to biases from computational procedures or biases in human interpretation due to the generally low resolution of these maps (see, e.g., experimental model clashscores higher than 10 in **Table 1**, which typically arise from fitting errors). In particular, for RNA, when the cryo-EM map has resolution worse than ∼3 Å, the separation between bases cannot be resolved and thus base placement can be highly dependent on the modeling approach used by the experimentalists. We therefore sought to rank CASP predictions based not on comparison to the reference coordinates provided by the experimenters (‘model-to-model’) but by comparison directly to the experimental maps (‘map-to-model’). The feasibility of refining these predictions to model the cryo-EM maps is discussed elsewhere in this issue^21^.

For all 6 RNA-only cryo-EM targets, there were models that could visually fit well into the maps (**Supplemental Figure 3**). To determine a quantitative ranking of predictor groups, previously available map-to-model metrics were computed (Methods; **Supplemental Figure 4**). These map-to-model metrics were developed to assess goodness of fit for models prepared with knowledge of maps; many were not designed to account for very poorly fitted models, with unmodeled density and atoms outside density, as we have here. For example, atomic inclusion^44^ penalizes predicted atoms that appear outside of density, and correlation coefficient at peaks (CC_peaks_)^37^ penalizes density that is not accounted for by a prediction. We attempted to find a combination of scores to balance these problems; however, in the end, we decided that no weighted combination of metrics was sufficient to enable ranking of all available models and predictors. Although overall correlation of map-to-model metrics to model-to-model metrics was high (**Supplemental Figure 5**), there were outliers receiving high map scores for poor models by, for example, condensing all atoms into a single small area, most notably group 238 (**Supplemental Figure 6C**). Thus, as in previous CASP evaluation for cryo-EM of protein targets^65^, we used a filter (**Supplemental Figure 6B**), only ranking models that exhibited sufficiently high model-to-model scores. Due to the size dependence of TM and GDT_TS noted above, we decided to set this cutoff based on RMSD. The correlation between metrics was generally improved after this filtering (**Supplemental Figure 6A** and **5B**).

For ranking, we selected a set of metrics that correlated well with visual inspections of fit and chose the standard measures of cross-correlation, accounting for modeled (CC_mask_) and unmodeled regions (CC_peaks_), and scores developed or shown to be most discriminatory for medium-resolution maps, atomic inclusion (AI), mutual information (MI), and Segment based Manders’ Overlap Coefficient (SMOC)^66,67^. We note that no metrics tested were RNA specific and can be used to assess any macromolecular complex. We used Z-score-based ranking, previously described, with uniform weight of the selected metrics:

AIchemy_RNA2 (TS232) achieved the highest Z_EM_ score, followed by Chen (TS287), GeneSilico (TS128), and RNApolis (TS081), and then others (**Figure 6A**). This ranking matched with the model-to-model assessment (orange bars in **Figure 6A**). This overall ranking was also maintained, barring group 238, without filtering out poor models (**Supplemental Figure 5A**); however, the filter should be maintained until Z_EM_ is robust to the problematic high scores of condensed models, by for example the inclusion of clashscore.

**FIGURE 6.**
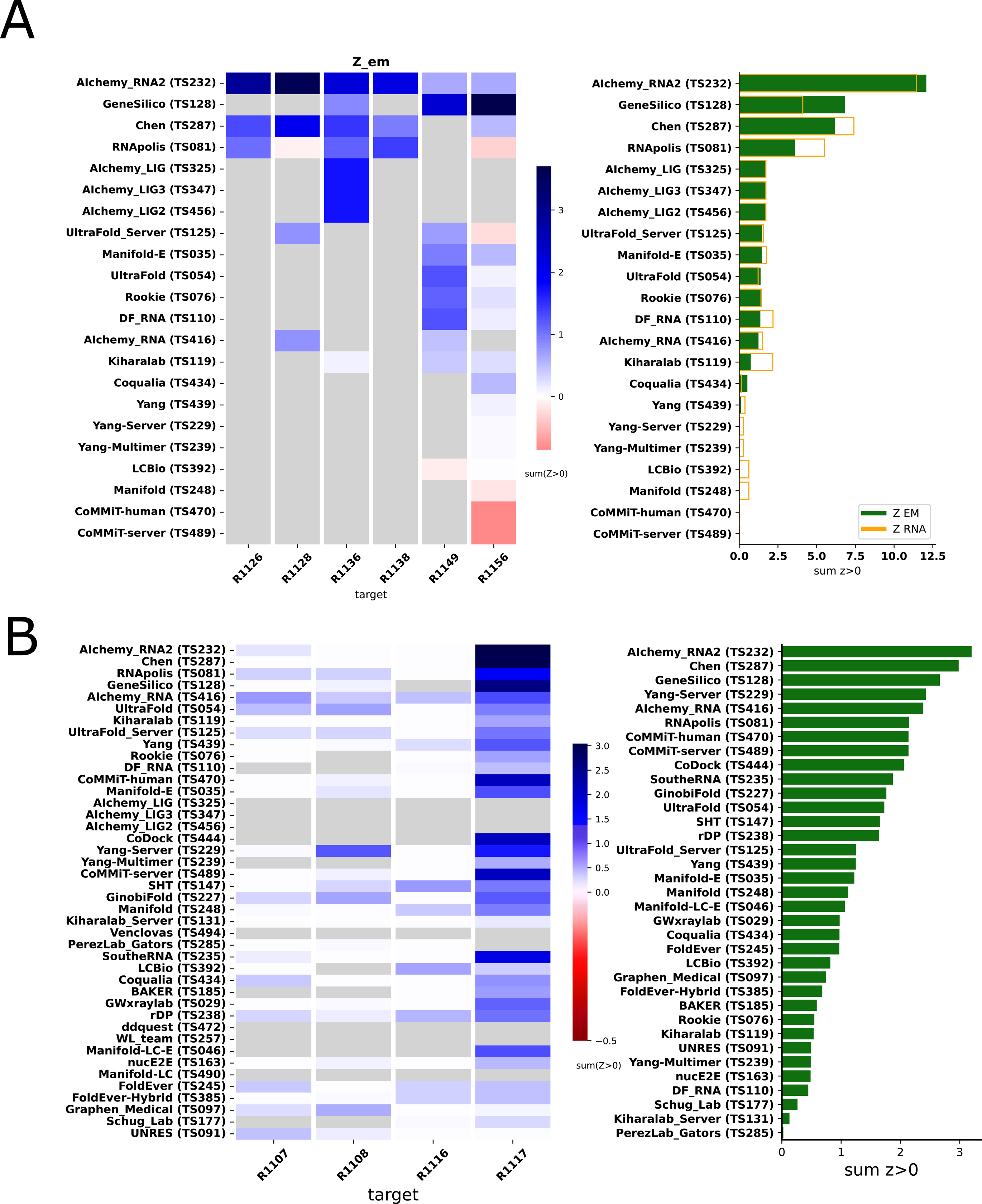
Ranking of CASP RNA predictions based on direct comparison to experimental data. (A) Ranking of six RNA-only cryo-EM targets based on Z-scores for map- to-model metrics (Z_EM_). Only a subset of models with clear alignments to maps were included in the comparison; see **Supplemental** Figure 5 for analysis over all models. **(B)** Group ranking for X-ray crystal structure targets based on Z-scores for metrics that directly compare the models to the crystallographic data (Z_MX_).

Overall, the results show that assessing models based on direct comparison to cryo-EM maps, appears feasible and that results are consistent with rankings based on model-to-model comparisons. Direct map-to-model assessments may be particularly important in future CASP events as prediction accuracy increases and approaches the level of detail obtained at typical cryo-EM map resolutions.

### 3.6 Ranking based on direct comparison to crystallographic data

In analogy to the map-based assessment of cryo-EM targets in the previous section, we investigated whether similar comparisons to the experimental data might enable ranking of the four RNA targets solved by X-ray macromolecular crystallography (MX). Similar to above, the only use of the experimentally derived model was to align predictor models. All predictor models were compared directly to the crystallographic data by first ideally placing the model using RNAalign^29^ and then calculating a Log Likelihood Gain (LLG) and translation-function Z-score (TFZ) with Phaser’s RNP search^46^ and a global map CC with phenix.get_cc_mtz_pdb^42^. We used a Z-based ranking after a round of outlier removal (see Methods) with a uniform weighting of these metrics:

The rankings are most strongly influenced by performance on R1117 since Z_MX_ scores for the other targets were relatively uniform and comparatively poor (**Figure 6B**). The top-ranking groups by this metric were TS232, TS287, and TS128 (AIchemy_RNA2, Chen, and GeneSilico, respectively), which were also the 3 groups that succeeded in follow up molecular replacement trials for R1117; see **Section 3.8** below.

### 3.7 CASP15 RNA models with accurate global folds miss detailed features and aspects of conformational heterogeneity

Ranking CASP15 RNA predictions based on the quantitative comparisons above highlighted several models for more detailed visual inspection, which revealed their potential and limitations. One example, the chimpanzee CPEB3 ribozyme R1108 (Figure 7), illustrates the use of the Deformation Profile and variable accuracy in targets of “medium” difficulty (**Table 1**). In **Figure 7A**, the superimposition of the experimental structure with the best model (TS232_4, from AIChemy_RNA2) is shown with the large deviations at the apical loops. The positions of these loops on the Deformation Profile (**Figure 7A-B**) are indicated highlighting the restricted regions with high discrepancies.

**FIGURE 7.**
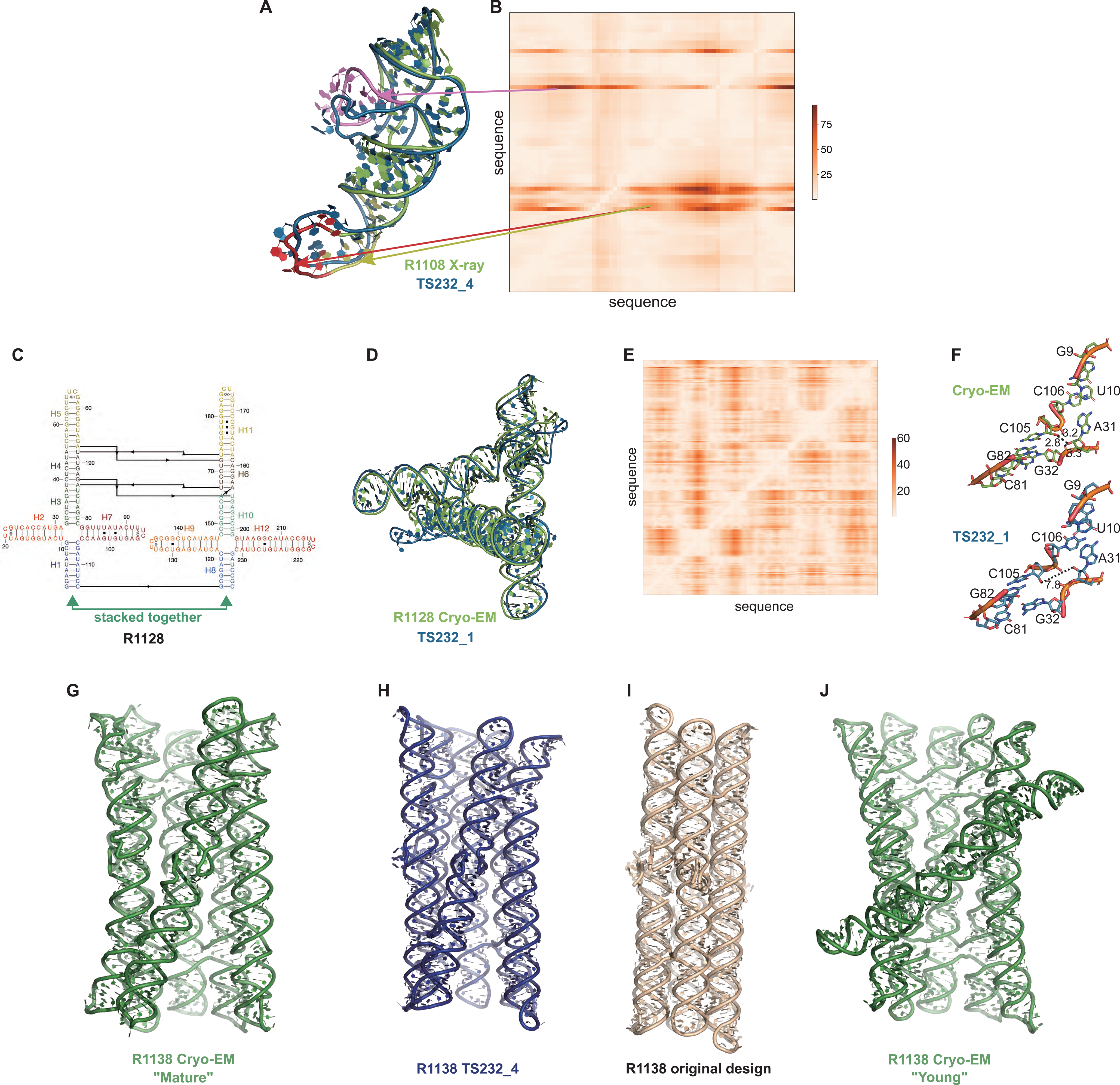
Detailed inspection of “medium” and “non-natural” targets. (A) For R1108 (chimpanzee CPEB3 ribozyme), superimposition of the experimental structure (green) with the best model (TS232_4 from AIChemy_RNA2, as blue, RMSD 4.5 Å) is shown. Notice the large deviations at the apical loops (as red, yellow and pink) and their positions on **(B)**, the Deformation Profile. **(C)** Diagram of the secondary structure (2D) of target R1128, a designed paranemic crossover triangle. The helices are numbered from H1 to H11. The secondary structure contains four 4-way junctions. In the two 4-way junctions drawn as “open”, helix H1 stacks with H2 and H3 with H7 for one 4-way junction and, for the second one, helix H8 stacks with H9 and H10 with H12. Helices H1 and H8 are stacked together. The pairs between G and U are marked by a dark dot (G•U pair). The Leontis-Westhof^81^ symbols are used to annotate the Watson-Crick/Sugar edge pair between G and U in the capping apical 5’UUCG3’ tetraloops. **(D)** Experimental structure (green) superimposed on the model TS232_1 (blue) with the lowest RMSD (4.3 Å). **(E)** The deformation profile (see Methods) between the same set of structures (at the right, the color scale where white represents excellent superimposition). The reddish regions indicate where the discrepancies are largest; they concentrate at the 4-way junctions where the experimental structure is more compact and with H-bonding contacts between the strands than the model structure as shown in **(F)**. **(G-J)** Models for R1128 (Paranemic Crossover Triangle, PXT). Cryo-EM of mature conformation **(G)** agrees better with blind CASP model TS232_4 **(H)** than with original models prepared by this nanostructure’s designers **(I)**. Cryo-EM also captured an early folding intermediate **(J)** that was not predicted well by any CASP15 groups.

One of the highly successful models is that of the paranemic crossover triangle (PTX) R1128, a molecule with no natural homologs whose difficulty for modeling was unclear before the CASP15 results^68^. It is a designed sequence made of four 4-way junctions and a co-axial stack between terminal helices (**Figure 7C-F**). The modeling success can be partly explained by the folding constraints of the design and the use of known structural modules. The helices are regular with known GU pairs and capping UNCG loops, without unpaired or bulging residues (**Figure 7C**). The tight junctions and the bulky RNA helices impose strong constraints on the fold and prevent knot formation (**Figure 7D**). The good accuracy of the modeling (TS232_1) with an RMSD of 4.3 Å and an INF of 0.88 is apparent in the deformation profile with a rather uniform deformation throughout (**Figure 7E**). The origins of the main errors are in the twist angles between stacked helices in the 4-way junctions that propagate maximally towards the apical loops (**Figure 7F**). In the experimentally determined structure, at those 4-way junctions, there are H-bonds linking one hydroxyl O2’ atom to an anionic phosphate oxygen of a residue on the crossing strand, maintaining a tight packing. These H-bonds are not present in the modeled structure, leading to a looser packing and slightly larger twist angle (**Figure 7G**). Despite these errors in fine details, the CASP15 blind model TS232_1 was closer to the cryo-EM-derived structure than the original model of the PTX structure designed by Andersen and colleagues (see paper co-submitted to CASP15 special issue^20^).

Indeed, for all four non-natural RNA targets in CASP15 (**Table 1**), the AIchemy_RNA2 group (TS232) submitted models that were visually accurate (**Figure 1**). Furthermore, this group, along with Chen (TS287) and RNAPolis (TS081) were notably separated from other groups, including all automated servers, for these non-natural targets, suggesting that these predictors benefitted from their human intuition to recognize the secondary structures and overall tertiary folds intended by the nanostructures’ human designers. Interestingly, in all four cases, the predictor groups were able to blindly predict structures that agreed better with the cryo-EM maps than the original models made by Andersen and colleagues when they designed the nanostructures. As another example, for R1138 (six-helix bundle, **Figure 7G-H**), the original design and the cryo-EM structure of the ‘mature’ form of the RNA agree in overall global fold, as reflected by a TM-score of 0.623, well above the 0.45 threshold (**Figure 7G-H)**. Nevertheless, the AIchemy_RNA2 model TS232_4 achieves an even higher TM-score of 0.800 (**Figure 7I**). These results suggest that, despite the lack of natural sequence homologs, “non-natural” RNA targets could be considered “easy” for 3D RNA structure prediction, as long as they are composed of readily identifiable helices and non-canonical motifs.

Interestingly, for the same R1138 six-helix-bundle, cryo-EM also captured a distinct ‘young’ structure for the RNA (**Figure 7J**) that is dominant immediately after the transcription of the RNA and requires hours to resolve into the ‘mature’ form^60^. The ‘young’ and ‘mature’ structures do not differ in their Watson-Crick-Franklin helices but, to interconvert, would require breaking of a kissing loop interaction, twisting of the two kissing elements about their helical axes, and then reformation of the kissing loop.^61^ None of the CASP models produced models close to the ‘young’ structure. Other natural and designed RNA systems are known to display similar kinetic traps and topological isomers^69,70^, and it will be interesting to see if in future CASPs, such conformations can be blindly predicted.

A common theme was that the model ordering as submitted by the predictor groups generally did not correspond to the ranking based on RMSDs (or other metrics) between experimental and model structures. This was the case for the R1108 and R1138 targets noted above, where the fourth models from group TS232, and not the first models, were most accurate. Overall, in 63% of the sets of CASP15 predictor submissions across all 12 RNA targets, a model submitted as 2-5 was better than model 1 by GDT_TS, and the difference in GDT_TS between model 1 and the top scoring model for each group was no lower than if model 1 had been randomly selected (**Supplemental Figure 7**). The models from group TS110 (DF_RNA) for the “difficult” target R1149 (the SL5 domain from SARS-CoV-2) provides an additional example. The best RMSD of all CASP15 submissions is model #2 by TS110 as depicted in **Figures 8A-D**. The RMSD between the experimental structure and TS110_2 is 6.9 Å (superposition shown on **Figure 8A** with the respective Deformation profile on **Figure 8B**). On the other hand, the RMSD between the experimental structure and first model TS110_1 is 21.7 Å. The superposition (**Figure 8C**) and the corresponding Deformation profile (**Figure 8D**) confirm that the global fold of TS110_1 is inaccurate despite its submission as model 1. In particular, the reddish regions indicate where the discrepancies are largest; they concentrate at the 4-way junctions where the experimental structure is more compact and with H-bonding contacts between the strands than the model structure as shown in **Figures** 8**A** and **7C**.

**FIGURE 8.**
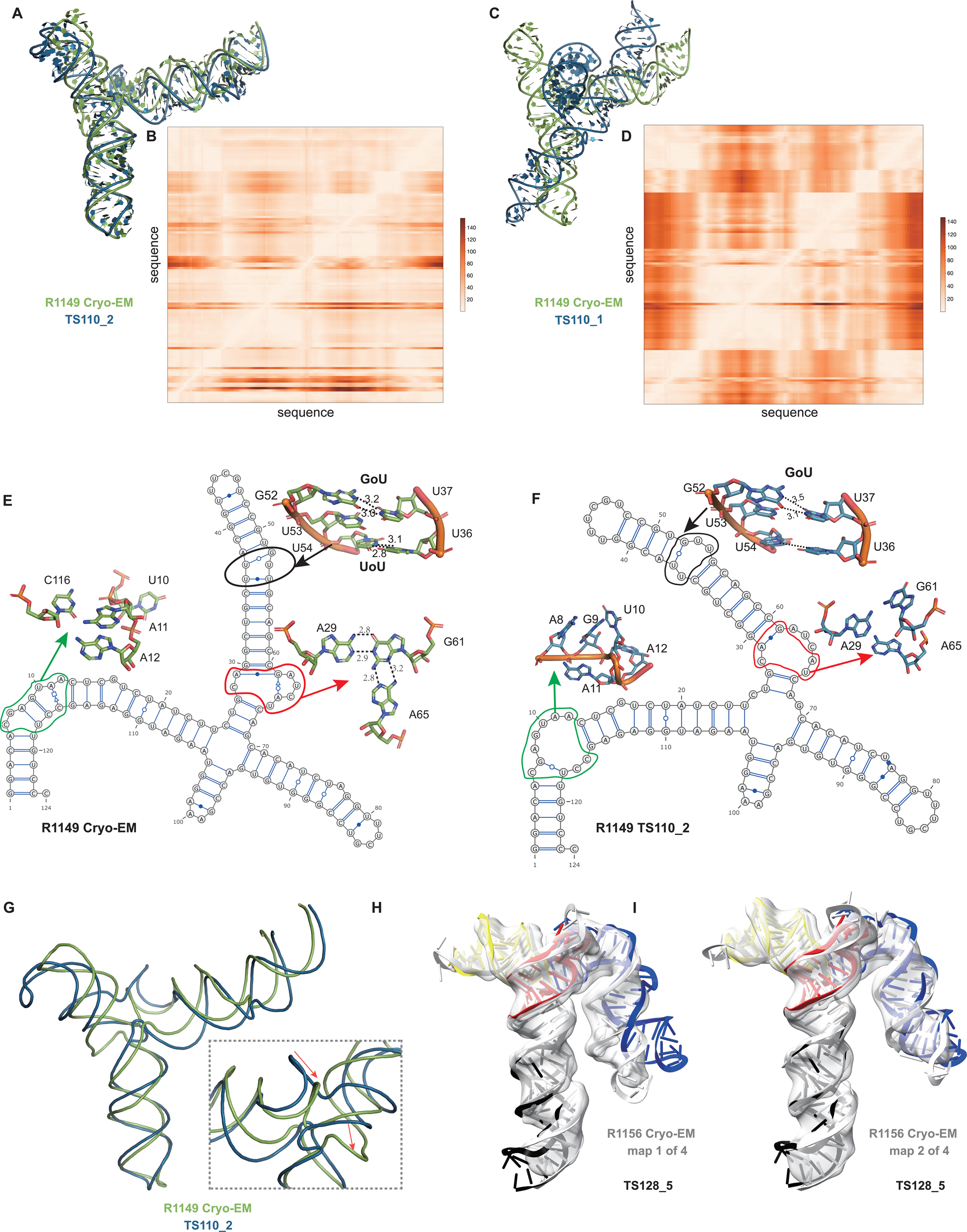
Detailed inspection of “difficult” targets, two coronavirus SL5 domains solved by cryo-EM. (A) Superposition between R1149 cryo-EM structure (first of 10 models representing experimental uncertainty) and the closest CASP15 prediction according to RMSD (TS110_2 with 6.9 Å). **(B)** Deformation profile between the same two structures. **(C)** Superposition between the experimental (R1149) and the model ranked #1 by the modeling group (TS110_1 with 21.7 Å). **(D)** Deformation profile between the same two structures. **(E)** Diagram of the secondary structure (2D) of target R1149 (first of 10 models representing experimental uncertainty). **(F)** Diagram of the secondary structure (2D) of the closest model TS110_2. The outlines indicate regions with large discrepancies due to wrong 2D pairs and absence of 3D pairs. For example, in the model structure, the U54/U36 pair is not present, and the region circled in green shows a region with high clashscore. **(G)** Backbone traces of the experimental (green) and model (blue) structures showing the overall fit of the helices; however, as shown in inset, the wrong choices in internal loops lead to large deviations in the path of the backbone at the central 4-way junction. **(H-I)** Experimental maps and models (gray) for R1156, whose cryo-EM data were subclassified into four separate conformations; conformation 1 **(H)** and 2 **(I)** compared to top scoring CASP prediction TS128_5 (color).

Further inspection of TS110_2 helps illustrate the requirement of paying attention to the non-Watson-Crick pairs beyond the standard Watson-Crick pairs of the secondary structure, both in prediction and in assessment of RNA targets resolved by cryo-EM. **Figures** 8**E** and 8**F** show the 2D structures for R1149 as derived from the cryo-EM map (**Figure 8E**) and the best RMSD model TS110_2 (**Figure 8F**) structures. The region within the black ellipse (**Figure 8G**) contains a GU and a UU pair, but in the modeled structure, only the GU pair is reproduced and, while the right Us face each other, they do not form a pair (**Figure 8H**). In the region circled in red, the fold of the single-stranded loop is missed and in the one circled in green, the fold leads to several bad contacts between residues, which may explain the rather high clashscore of 31 for TS110_2, despite the overall good fit in the relative orientations between the helices (**Figure 8A**). It is important to note that for these regions, alternative structures in the experimentalists’ 10-model cryo-EM ensemble show breaking of the features, similar to the prediction TS110_2; and so it is possible that the conformations modeled in TS110_2 occur in the actual cryo- ensemble for the target R1149. Nevertheless, these model discrepancies lead to deviations of the strands in the four-way junction that, in turn, lead to variations in the arms at the junction (**Figure 8G**). Indeed, all 10 members of the experimental cryo-EM ensemble show complete base pairing at the molecule’s central four-way junction, which is inconsistent with incomplete junction base pairing in TS110_2 (**Figure 8F**).

The presence of alternative structures, noted above for the non-natural six helix bundle R1138, was a common theme in RNA targets in CASP (**Table 1**), and was particularly interesting in one target with continuous heterogeneity. R1156 is a homolog of the same SARS-CoV-2 SL5 domain as R1149, and showed flexibility in one helix (blue, **Figure 8H-I**), which was represented in the cryo-EM analysis as four sub-classified maps. Comparing models directly to these experimental maps highlighted models of particular excellent quality that fit into the maps nearly as well as the reference models prepared by experimentalists using the maps (**Supplemental Figure 3**). In particular, the model TS128_5 from GeneSilico fits into the experimental map with excellent scores (**Figure 8H-I**). Fitting this model into the highest resolution of these four maps, conformation 1, we can see visually and numerically, that the model fits well with respect to 3 helices but poorly with respect to the flexible helix (**Figure 8H**). However, the model fits better in the second conformation, obtaining map-to-model atomic inclusion scores comparable to scores achieved by models derived with knowledge of the map (**Figure 8I**). This comparison revealed the importance of representing the ensemble of structures the RNA can form so as to not penalize prediction of structures that do form but can not be captured by a single experimental structure.

In summary, inspection of top-ranked CASP15 RNA models confirms, in each case, good prediction of global fold but also reveals fine details and/or aspects of conformational heterogeneity that have not been captured by the models. Ordering each set of five models by the predictor groups also typically did not correlate with the models’ accuracy. Similar conclusions for R1108, R1128, R1138, R1149 and R1156 based on alternative analyses by RNA experimental groups, are described in a separate paper prepared for the CASP issue.^20^

### 3.8 Potential utility of RNA models for molecular replacement

The general global fold accuracy of the CASP15 RNA tertiary structure models motivated us to explore their potential utility for phasing X-ray diffraction data by molecular replacement, which has previously been carried out in very few cases^71^. While we began these explorations in studies described above to rank models based on agreement with X-ray data (**Section 3.6**), such scores based on optimal placements do not necessarily reflect models’ value as search models for real-world Molecular Replacement (MR). For example, a largely accurate model may prove unsuccessful if inaccurately modeled portions lead to severe crystal lattice packing clashes.

We therefore carried out more realistic MR runs, first, on all unmodified models of R1117. This initial analysis was restricted to R1117 since visual examination and LLG calculations suggested that models of other targets would require some kind of editing to succeed (see next). Across the up to five models submitted by groups, we found that 3 out of 34 groups succeeded with at least one model, using global map correlation coefficient CC > 0.2 as the criterion of success (**Supplemental Figure 8**). Among these successful groups, however, the quality of MR solutions varied significantly. The highest LLG was 110 for model 128_2 but results in a poor R_free_ of 54% after refinements in Refmac5; in contrast, the lowest R_free_ was 39% for model 287_3 from the Chen group after refinement with Refmac5^72^ despite this model giving a worse LLG in the MR trials. **Figure 9** shows the successful solution with model 287_3.

**FIGURE 9.**
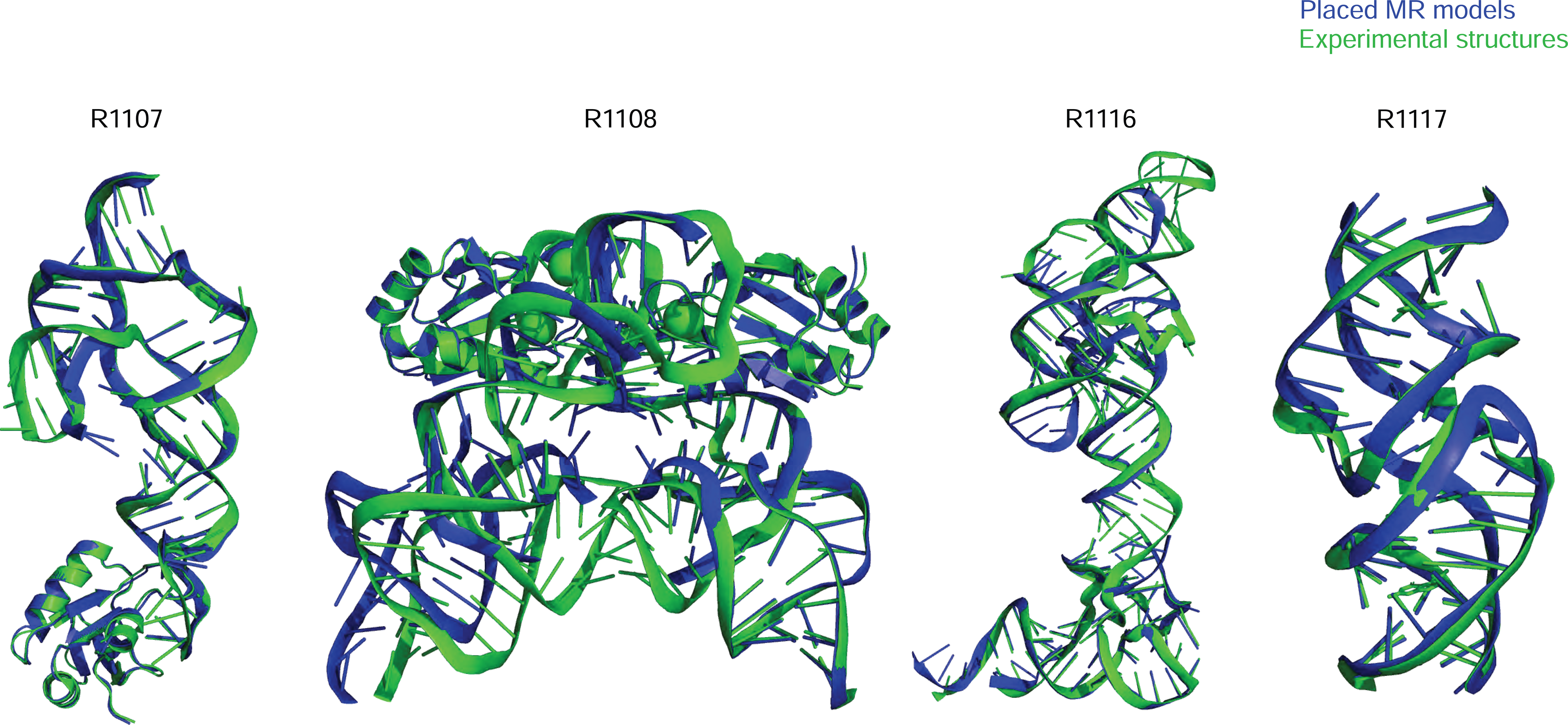
Molecular replacement (MR) of X-ray crystallographic data using CASP15 models (and AlphaFold 2 models of U1ABD in the cases of R1107 and R1108). Group TS232 models formed the basis of all successful search models shown except R1117 (group TS287).

For the other three CASP targets solved by X-ray diffraction, visual inspection and the Z_MX_ values in **Figure 6B** made clear that editing of the predictions would be required for successful MR, and, to focus resources, models from TS232 (AIchemy_RNA2) were subjected to various editing procedures. For solution of the two CPEB3 ribozymes, R1107 (one protein chain, one RNA chain) and R1108 (two protein chains, two RNA chains), the structural variance observed in group TS232 models after structural alignment with Theseus^73^ was used as an indication of local prediction reliability and divergent regions removed before the edited model_1 was used as a search model. This approach borrows from that taken for proteins by the MR pipeline AMPLE^74^. R1107 was successfully solved by first placing the protein chain then the edited RNA search model, both with Phaser. The result (**Figure 9**) has an R_free_ of 26% and visible density for the missing part of the RNA molecule confirms that it could be readily refined and completed. R1108, a close homolog of R1107, proved much more difficult to solve, perhaps owing to the different conformations observed between the two RNA chains in the asymmetric unit. When attempting to solve this structure similarly (protein first then RNA) we could place the protein component, but the RNA component was reversed, providing only a partial solution. The truncated group TS232 models for R1108 were of a sufficient quality to solve R1107 and the resulting protein/RNA complex could then be used to solve R1108 with an R_free_ of 41%.

Inspection of the Group 232 models for R1116 showed that more extensive model editing would be required. A modified version of Slice’N’Dice^53^ was therefore used to split model_1 into three structural units. A portion comprising nucleotides 1-24;125-157 could then be placed with MOLREP which indicated a partial solution after refinement (R_fact_ 48%, R_free_ 52%). Three copies of a second fragment comprising 38-63 could then be placed to largely complete the structure with Phaser scores of LLG: 1324 and TFZ: 9.6. These values are unambiguously indicative of successful Molecular Replacement: for example, TFZ > 8 corresponds to ‘Definitely’ solved according to Phaser software guidance.^75^ The result (**Figure 9**) has an R_free_ of 43%, an acceptable value for a model immediately after MR. These results demonstrate that all of the RNA crystal structure targets in CASP15 could, one way or another, be solved by MR, although it is recognised that further refinement and completion (not attempted here) could be challenging, especially at 3.0 Å or worse resolution.

## 4 Discussion

CASP15 enabled a timely assessment of 3D RNA structure prediction, with 8 RNA targets solved by cryo-EM and 4 by X-ray crystallography. Forty two predictors from 25 research centers made submissions for at least one of these targets, many of whom had not published studies on RNA prior to CASP but explored deep learning approaches that were novel for the RNA field. The twelve RNA targets ranged in difficulty from “easy”, with clearly identifiable templates in the structure database, to “difficult”, with no templates. When looking at all five submissions for each target, visually good predictions were submitted for all 10 RNA-only targets, including 4 non-natural RNA targets that had no global homology to previously solved structures. Two protein-RNA complexes were not modeled accurately.

Quantitative rankings of predictor groups were carried out by independent teams, based on RMSD and INF metrics developed in the RNA-Puzzles trials and based on TM-score, GDT_TS, and lDDT more familiar to protein structure assessments in prior CASP. Both rankings agreed in placing TS232 (AIchemy_RNA2) first, TS287 (Chen) second, and TS128 and TS081 (GeneSilico and RNAPolis) as tied for third. These rankings were also confirmed by analyses comparing predicted models to maps (for cryo-EM targets) and statistics related to molecular replacement (for X-ray crystal targets). The top-ranked models for the 10 RNA-only targets captured global folds well, as assessed by visual inspection and by achievement of GDT_TS values greater than 45 and/or TM-score values greater than 0.45. Nevertheless, fine details such as non canonical pairs and hydrogen bonding at junctions were inaccurate in these models, even when taking into account sources of uncertainty for the experimental structures. Conformational heterogeneity in some targets, R1136 and R1156, was indicated by the presence of multiple structures captured by cryo-EM but was not captured by any group in their range of submitted models (**Supplemental Figure 9**). Despite these caveats, the general global fold accuracy for RNA-only targets — even those without homologs of known structure — and the ability of models, with some curation, to enable molecular replacement of all 4 X-ray diffraction data sets suggests reason for optimism.

Has there been improvement in RNA modeling in CASP15 compared to prior RNA-puzzles? Achieving accurate positioning of helices with respect to each other by modeling is often feasible when the single-stranded segments are short and unpaired because RNA helices are bulky and the interconnecting strands each have a polarity, leading to a reduced search space for modeling helix arrangements. The good helix positioning observed here in CASP15 was also regularly observed during previous RNA-Puzzles assessments and in previous RNA modeling efforts. During this CASP15 experiment, some research groups tried to use prediction approaches that were similar to AI-based methods for predicting the structure of proteins. For example, AlChemy_RNA^76^ uses an end-to-end differentiable network inspired by AlphaFold 2^51^. However, these AI-based predictions did not perform as well as expected and did not surpass prediction methods previously tested in RNA-Puzzles (SimRNA, Chen, RNApolis), which have been continuously improving for the past decade. The AI-based approaches^76,77^ also failed to demonstrate the accuracy claimed in their preprint papers, perhaps due to the limited amount of training data.

In addition to not using deep learning, the top four RNA predictors shared the property that they were not servers and, based on their own accounts (see papers co-submitted for this CASP issue^64^), they appeared to still make use of human intuition. While there were cases where server models were more accurate than ‘human’ models from the same laboratory (e.g., Yang), generally server models were worse in quality than the top 4 human predictor groups. Going forward, an important frontier for the RNA structure prediction field to focus on will be automation, so that methods can be more widely used and applied at the genomic scale, as is now the case for protein structure prediction methods. While the sparser data available for RNA structure, compared to protein structure, has complicated development of robust deep learning algorithms, recent accelerations in RNA structure determination – particularly from cryo-EM^12^ – and the availability of high-throughput sequencing-based methods sensitive to RNA structure^78^ may help close the gap between RNA and protein computational methods. Interestingly, secondary structures from even the top server predictions were poorer than those from ‘human’ groups, highlighting an area of potentially immediate improvement.

In addition to being the first CASP experiment for RNA structure prediction, CASP15 was also the first CASP experiment for RNA structure assessment, and future CASP RNA trials can benefit from some lessons learned by the assessors, three of which we discuss here. First, CASP15 included few truly difficult RNA targets, and these were solved by cryo-EM at resolutions worse than 3 Å. It will be important for upcoming CASP competitions to bring in experimental groups solving natural RNA targets without previously solved homologs at near-atomic resolution. Such molecules are being discovered and structurally characterized at increasing frequency, particularly for biologically interesting RNA-protein complexes. It may also be useful to develop a fully automated classification scheme for easy, medium, and difficult RNA targets and separately assess targets from these categories, as was traditional in CASP before the success of deep learning approaches rendered these categories less useful for proteins.

Second, while only 2 of 12 targets in CASP15 were RNA-protein complexes, it seems feasible that CASP16 will involve more RNA-protein complexes, given their biological importance and amenability to cryo-EM. For assessment, it will therefore be increasingly important to develop quantitative metrics that make sense across RNA, protein, and RNA-protein interfaces. We found here that standard metrics for protein structure accuracy assessment, GDT_TS and TM- score, were useful in ranking RNA models, but their values for visually excellent RNA models seemed anomalously low for large and small targets, respectively. More local measures of accuracy, like lDDT or assessments of contact accuracy, appeared useful here for both RNA and RNA-protein targets. These more local measures may be less affected by length variation and also more robust to dynamic fluctuations that appear common in large, extended RNA structures. The recent availability of lDDT for RNA may allow more testing of this metric in continuous trials like CAMEO and RNA-Puzzles before the next CASP.

Third, many and perhaps most of the CASP15 RNA targets showed conformational flexibility, e.g., as evidenced by differences in conformations of different monomers in crystallographic asymmetric units or, in cryo-ensembles captured by electron microscopy as classes of conformations separable by automated subclassification and/or 3D variability analysis^39^. In the current assessment, predictor groups were scored based on the best observed agreement of all their submitted models vs. all available experimental models, effectively assuming that modelers were predicting single structures. Modeling of the full ensemble nature of these RNA systems was neither incentivized nor assessed. In future CASPs, acceptance of multi-model ensembles (with e.g., 100’s or 1000’s of models within each of 5 ensembles), rather than separate single-structure models, would better incentivize development of methods for predicting conformational ensembles of macromolecules, including molecular dynamics methods that have been previously difficult to assess. Furthermore, scoring of these ensembles directly against data should be feasible; e.g., log-likelihood frameworks and GPU-enabled software^79^ might enable predicted multi-model cryo-ensembles to be compared to the entire collection of electron micrographs collected for a target.

## Acknowledgements

We thank Ebbe Andersen for sharing in-house models of RNA designs; Nick Grishin and Lisa Kinch for advice on Z-score computations; Marta Szachniuk and Maciej Antczak for advice and numerical cross-checks for RNA assessment metrics^80^; Gabriel Studer for extending lDDT to RNA; Marcin Magnus for advice on RNA model cleanup and assessment tools; Adam Zemla for extending LGA to RNA; Chengxin Zhang for advice on TM-score calculations; the RNA-puzzles modeling community for their participation as predictors; all experimentalists who contributed the RNA structures; and Andriy Kryshtafovych, Krzystof Fidelis, and John Moult for the invitation and dedicated support to bring RNA model assessment to CASP. This research was supported by Stanford Bio-X (to R.D. and R.C.K.); Stanford Gerald J. Lieberman Fellowship (to R.R.); the National Institutes of Health (R35 GM122579 to R.D.), the Howard Hughes Medical Institute (HHMI, to R.D.); Biotechnology and Biological Sciences Research Council (BBSRC) grant BB/S007105/1 (D.J.R.); Leibniz ScienceCampus InterACt, funded by the BWFGB Hamburg and the Leibniz Association (M.T.); the Natural Science Foundation of China (32270707 to Z.M.), the National Key R&D Program of China (2021YFF1200900, 2021YFF1200903 to Z.M.); R&D Program of Guangzhou Laboratory (Grant No. SRPG22-003, SRPG22-006, SRPG22-007 to Z.M.); French National Research Agency (LABEX: ANR-10-LABX-0036_NETRNA, Investments for the Future program, to E.W.). This article is subject to HHMI’s Open Access to Publications policy. HHMI lab heads have previously granted a nonexclusive CC BY 4.0 license to the public and a sublicensable license to HHMI in their research articles. Pursuant to those licenses, the author-accepted manuscript of this article can be made freely available under a CC BY 4.0 license immediately upon publication.

## Author contributions

R.D., R.C.K., and E.W. coordinated the research. Z.M. and E.W. and, independently, R.D., R.C.K., P.P., and R.R. designed and conducted global model-to-model RNA assessments. T.M. and M.T. designed and conducted protein-RNA folding pattern assessment. R.C.K. designed and conducted cryo-EM map-to-model assessment. A.J.S., R.M.K., and D.J.R. designed and conducted molecular replacement tests. R.D., R.C.K., A.J.S., T.M., P.P., R.R., and E.W. drafted the paper and prepared figures. F.B. prepared figures; and Z.M., P.P., F.B., and R.C.K. designed and curated code repositories.

## Conflict of Interest

All authors declare that they have no competing interests.

## Notes

https://github.com/DasLab/casp-rna

https://predictioncenter.org/casp15/results.cgi?view=targets&tr_type=rna

### Competing Interest Statement

The authors have declared no competing interest.

### Summary of Updates

- Supplemental Information Figures added. - Figure 4A row order corrected. - Minor typos corrected throughout.

## References

1. Holley, R. W., Apgar, J., Everett, G. A., Madison, J. T., Marquisee, M., Merrill, S. H., Penswick, J. R. & Zamir, A. STRUCTURE OF A RIBONUCLEIC ACID. Science 147, 1462– 1465 (1965).

2. Madison, J. T., Everett, G. A. & Kung, H. Nucleotide sequence of a yeast tyrosine transfer RNA. Science 153, 531–534 (1966).

3. Fuller, W. & Hodgson, A. Conformation of the anticodon loop intRNA. Nature 215, 817–821 (1967).

4. Levitt, M. Detailed molecular model for transfer ribonucleic acid. Nature 224, 759–763 (1969).

5. Hingerty, B., Brown, R. S. & Jack, A. Further refinement of the structure of yeast tRNAPhe. J. Mol. Biol. 124, 523–534 (1978).

6. Sussman, J. L., Holbrook, S. R., Warrant, R. W., Church, G. M. & Kim, S. H. Crystal structure of yeast phenylalanine transfer RNA. I. Crystallographic refinement. J. Mol. Biol. 123, 607–630 (1978).

7. Westhof, E. & Leontis, N. B. An RNA-centric historical narrative around the Protein Data Bank. J. Biol. Chem. 296, 100555 (2021).

8. Cruz, J. A., Blanchet, M.-F., Boniecki, M., Bujnicki, J. M., Chen, S.-J., Cao, S., Das, R., Ding, F., Dokholyan, N. V., Flores, S. C., Huang, L., Lavender, C. A., Lisi, V., Major, F., Mikolajczak, K., Patel, D. J., Philips, A., Puton, T., Santalucia, J., Sijenyi, F., Hermann, T., Rother, K., Rother, M., Serganov, A., Skorupski, M., Soltysinski, T., Sripakdeevong, P., Tuszynska, I., Weeks, K. M., Waldsich, C., Wildauer, M., Leontis, N. B. & Westhof, E. RNA-Puzzles: a CASP-like evaluation of RNA three-dimensional structure prediction. RNA 18, 610–625 (2012).

9. Miao, Z., Adamiak, R. W., Antczak, M., Batey, R. T., Becka, A. J., Biesiada, M., Boniecki, M. J., Bujnicki, J. M., Chen, S.-J., Cheng, C. Y., Chou, F.-C., Ferré-D’Amaré, A. R., Das, R., Dawson, W. K., Ding, F., Dokholyan, N. V., Dunin-Horkawicz, S., Geniesse, C., Kappel, K., Kladwang, W., Krokhotin, A., Łach, G. E., Major, F., Mann, T. H., Magnus, M., Pachulska-Wieczorek, K., Patel, D. J., Piccirilli, J. A., Popenda, M., Purzycka, K. J., Ren, A., Rice, G. M., Santalucia, J., Jr, Sarzynska, J., Szachniuk, M., Tandon, A., Trausch, J. J., Tian, S., Wang, J., Weeks, K. M., Williams, B., 2nd, Xiao, Y., Xu, X., Zhang, D., Zok, T. & Westhof, E. RNA-Puzzles Round III: 3D RNA structure prediction of five riboswitches and one ribozyme. RNA 23, 655–672 (2017).

10. Miao, Z., Adamiak, R. W., Antczak, M., Boniecki, M. J., Bujnicki, J., Chen, S.-J., Cheng, C. Y., Cheng, Y., Chou, F.-C., Das, R., Dokholyan, N. V., Ding, F., Geniesse, C., Jiang, Y., Joshi, A., Krokhotin, A., Magnus, M., Mailhot, O., Major, F., Mann, T. H., Piątkowski, P., Pluta, R., Popenda, M., Sarzynska, J., Sun, L., Szachniuk, M., Tian, S., Wang, J., Wang, J., Watkins, A. M., Wiedemann, J., Xiao, Y., Xu, X., Yesselman, J. D., Zhang, D., Zhang, Y., Zhang, Z., Zhao, C., Zhao, P., Zhou, Y., Zok, T., Żyła, A., Ren, A., Batey, R. T., Golden, B. L., Huang, L., Lilley, D. M., Liu, Y., Patel, D. J. & Westhof, E. RNA-Puzzles Round IV: 3D structure predictions of four ribozymes and two aptamers. RNA 26, 982–995 (2020).

11. Miao, Z., Adamiak, R. W., Blanchet, M.-F., Boniecki, M., Bujnicki, J. M., Chen, S.-J., Cheng, C., Chojnowski, G., Chou, F.-C., Cordero, P., Cruz, J. A., Ferré-D’Amaré, A. R., Das, R., Ding, F., Dokholyan, N. V., Dunin-Horkawicz, S., Kladwang, W., Krokhotin, A., Lach, G., Magnus, M., Major, F., Mann, T. H., Masquida, B., Matelska, D., Meyer, M., Peselis, A., Popenda, M., Purzycka, K. J., Serganov, A., Stasiewicz, J., Szachniuk, M., Tandon, A., Tian, S., Wang, J., Xiao, Y., Xu, X., Zhang, J., Zhao, P., Zok, T. & Westhof, E. RNA-Puzzles Round II: assessment of RNA structure prediction programs applied to three large RNA structures. RNA 21, 1066–1084 (2015).

12. Das, R. RNA structure: a renaissance begins? Nat. Methods 18, 439 (2021).

13. Pereira, J., Simpkin, A. J., Hartmann, M. D., Rigden, D. J., Keegan, R. M. & Lupas, A. N. High-accuracy protein structure prediction in CASP14. Proteins 89, 1687–1699 (2021).

14. Hogan, M. J. & Pardi, N. mRNA Vaccines in the COVID-19 Pandemic and Beyond. Annu. Rev. Med. 73, 17–39 (2022).

15. Lensink, M. F., Brysbaert, G., Mauri, T., Nadzirin, N., Velankar, S., Chaleil, R. A. G., Clarence, T., Bates, P. A., Kong, R., Liu, B., Yang, G., Liu, M., Shi, H., Lu, X., Chang, S., Roy, R. S., Quadir, F., Liu, J., Cheng, J., Antoniak, A., Czaplewski, C., Giełdoń, A., Kogut, M., Lipska, A. G., Liwo, A., Lubecka, E. A., Maszota-Zieleniak, M., Sieradzan, A. K., Ślusarz, R., Wesołowski, P. A., Zięba, K., Del Carpio Muñoz, C. A., Ichiishi, E., Harmalkar, A., Gray, J. J., Bonvin, A. M. J. J., Ambrosetti, F., Vargas Honorato, R., Jandova, Z., Jiménez-García, B., Koukos, P. I., Van Keulen, S., Van Noort, C. W., Réau, M., Roel-Touris, J., Kotelnikov, S., Padhorny, D., Porter, K. A., Alekseenko, A., Ignatov, M., Desta, I., Ashizawa, R., Sun, Z., Ghani, U., Hashemi, N., Vajda, S., Kozakov, D., Rosell, M., Rodríguez-Lumbreras, L. A., Fernandez-Recio, J., Karczynska, A., Grudinin, S., Yan, Y., Li, H., Lin, P., Huang, S.-Y., Christoffer, C., Terashi, G., Verburgt, J., Sarkar, D., Aderinwale, T., Wang, X., Kihara, D., Nakamura, T., Hanazono, Y., Gowthaman, R., Guest, J. D., Yin, R., Taherzadeh, G., Pierce, B. G., Barradas-Bautista, D., Cao, Z., Cavallo, L., Oliva, R., Sun, Y., Zhu, S., Shen, Y., Park, T., Woo, H., Yang, J., Kwon, S., Won, J., Seok, C., Kiyota, Y., Kobayashi, S., Harada, Y., Takeda-Shitaka, M., Kundrotas, P. J., Singh, A., Vakser, I. A., Dapkūnas, J., Olechnovič, K., Venclovas, Č., Duan, R., Qiu, L., Xu, X., Zhang, S., Zou, X. & Wodak, S. J. Prediction of protein assemblies, the next frontier: The CASP14-CAPRI experiment. Proteins 89, 1800–1823 (2021).

16. Lensink, M. F., Brysbaert, G., Nadzirin, N., Velankar, S., Chaleil, R. A. G., Gerguri, T., Bates, P. A., Laine, E., Carbone, A., Grudinin, S., Kong, R., Liu, R.-R., Xu, X.-M., Shi, H., Chang, S., Eisenstein, M., Karczynska, A., Czaplewski, C., Lubecka, E., Lipska, A., Krupa, P., Mozolewska, M., Golon, Ł., Samsonov, S., Liwo, A., Crivelli, S., Pagès, G., Karasikov, M., Kadukova, M., Yan, Y., Huang, S.-Y., Rosell, M., Rodríguez-Lumbreras, L. A., Romero-Durana, M., Díaz-Bueno, L., Fernandez-Recio, J., Christoffer, C., Terashi, G., Shin, W.-H., Aderinwale, T., Maddhuri Venkata Subraman, S. R., Kihara, D., Kozakov, D., Vajda, S., Porter, K., Padhorny, D., Desta, I., Beglov, D., Ignatov, M., Kotelnikov, S., Moal, I. H., Ritchie, D. W., Chauvot de Beauchêne, I., Maigret, B., Devignes, M.-D., Ruiz Echartea, M. E., Barradas-Bautista, D., Cao, Z., Cavallo, L., Oliva, R., Cao, Y., Shen, Y., Baek, M., Park, T., Woo, H., Seok, C., Braitbard, M., Bitton, L., Scheidman-Duhovny, D., Dapkūnas, J., Olechnovič, K., Venclovas, Č., Kundrotas, P. J., Belkin, S., Chakravarty, D., Badal, V. D., Vakser, I. A., Vreven, T., Vangaveti, S., Borrman, T., Weng, Z., Guest, J. D., Gowthaman, R., Pierce, B. G., Xu, X., Duan, R., Qiu, L., Hou, J., Ryan Merideth, B., Ma, Z., Cheng, J., Zou, X., Koukos, P. I., Roel-Touris, J., Ambrosetti, F., Geng, C., Schaarschmidt, J., Trellet, M. E., Melquiond, A. S. J., Xue, L., Jiménez-García, B., van Noort, C. W., Honorato, R. V., Bonvin, A. M. J. J. & Wodak, S. J. Blind prediction of homo- and hetero-protein complexes: The CASP13-CAPRI experiment. Proteins 87, 1200–1221 (2019).

17. Lensink, M. F., Velankar, S., Baek, M., Heo, L., Seok, C. & Wodak, S. J. The challenge of modeling protein assemblies: the CASP12-CAPRI experiment. Proteins 86 **Suppl 1**, 257– 273 (2018).

18. Lensink, M. F., Velankar, S., Kryshtafovych, A., Huang, S.-Y., Schneidman-Duhovny, D., Sali, A., Segura, J., Fernandez-Fuentes, N., Viswanath, S., Elber, R., Grudinin, S., Popov, P., Neveu, E., Lee, H., Baek, M., Park, S., Heo, L., Rie Lee, G., Seok, C., Qin, S., Zhou, H.- X., Ritchie, D. W., Maigret, B., Devignes, M.-D., Ghoorah, A., Torchala, M., Chaleil, R. A. G., Bates, P. A., Ben-Zeev, E., Eisenstein, M., Negi, S. S., Weng, Z., Vreven, T., Pierce, B. G., Borrman, T. M., Yu, J., Ochsenbein, F., Guerois, R., Vangone, A., Rodrigues, J. P. G. L.M., van Zundert, G., Nellen, M., Xue, L., Karaca, E., Melquiond, A. S. J., Visscher, K., Kastritis, P. L., Bonvin, A. M. J. J., Xu, X., Qiu, L., Yan, C., Li, J., Ma, Z., Cheng, J., Zou, X., Shen, Y., Peterson, L. X., Kim, H.-R., Roy, A., Han, X., Esquivel-Rodriguez, J., Kihara, D., Yu, X., Bruce, N. J., Fuller, J. C., Wade, R. C., Anishchenko, I., Kundrotas, P. J., Vakser, I. A., Imai, K., Yamada, K., Oda, T., Nakamura, T., Tomii, K., Pallara, C., Romero-Durana, M., Jiménez-García, B., Moal, I. H., Férnandez-Recio, J., Joung, J. Y., Kim, J. Y., Joo, K., Lee, J., Kozakov, D., Vajda, S., Mottarella, S., Hall, D. R., Beglov, D., Mamonov, A., Xia, B., Bohnuud, T., Del Carpio, C. A., Ichiishi, E., Marze, N., Kuroda, D., Roy Burman, S. S., Gray, J. J., Chermak, E., Cavallo, L., Oliva, R., Tovchigrechko, A. & Wodak, S. J. Prediction of homoprotein and heteroprotein complexes by protein docking and template-based modeling: A CASP-CAPRI experiment. Proteins 84 **Suppl 1**, 323–348 (2016).

19. Ozden, B., Kryshtafovych, A. & Karaca, E. Assessment of the CASP14 assembly predictions. Proteins 89, 1787–1799 (2021).

20. Kretsch, R. C., Andersen, E. S., Bujnicki, J. M., Chiu, W. & Das, R. RNA target highlights in CASP15: Evaluation of predicted models by structure providers. authorea.com doi:10.22541/au.168487314.47726735

21. Thomas Mulvaney, Rachael C. Kretsch, Luc Elliott, Joe Beton, Andriy Kryshtafovych, Daniel Rigden, Rhiju Das, Maya Topf. CASP15 cryoEM protein and RNA targets: refinement and analysis using experimental maps. Authorea preprint doi:https://www.authorea.com/doi/full/10.22541/au.168742148.85721558/v1

22. Parisien, M., Cruz, J. A., Westhof, E. & Major, F. New metrics for comparing and assessing discrepancies between RNA 3D structures and models. RNA 15, 1875–1885 (2009).

23. Gendron, P., Lemieux, S. & Major, F. Quantitative analysis of nucleic acid three-dimensional structures. J. Mol. Biol. 308, 919–936 (2001).

24. 24. Comparison of the predicted and observed secondary structure of T4 phage lysozyme. *Biochimica et Biophysica Acta (BBA) - Protein Structure* 405, 442–451 (1975).

25. Gorodkin, J., Stricklin, S. L. & Stormo, G. D. Discovering common stem–loop motifs in unaligned RNA sequences. Nucleic Acids Res. 29, 2135–2144 (2001).

26. Davis, I. W., Leaver-Fay, A., Chen, V. B., Block, J. N., Kapral, G. J., Wang, X., Murray, L. W., Arendall, W. B., 3rd, Snoeyink, J., Richardson, J. S. & Richardson, D. C. MolProbity: all-atom contacts and structure validation for proteins and nucleic acids. Nucleic Acids Res. 35, W375–83 (2007).

27. Murray, L. J. W., Arendall, W. B., 3rd, Richardson, D. C. & Richardson, J. S. RNA backbone is rotameric. Proc. Natl. Acad. Sci. U. S. A. 100, 13904–13909 (2003).

28. Word, J. M., Lovell, S. C., LaBean, T. H., Taylor, H. C., Zalis, M. E., Presley, B. K., Richardson, J. S. & Richardson, D. C. Visualizing and quantifying molecular goodness-of-fit: small-probe contact dots with explicit hydrogen atoms. J. Mol. Biol. 285, 1711–1733 (1999).

29. Gong, S., Zhang, C. & Zhang, Y. RNA-align: quick and accurate alignment of RNA 3D structures based on size-independent TM-scoreRNA. Bioinformatics 35, 4459–4461 (2019).

30. Zok, T., Popenda, M. & Szachniuk, M. MCQ4Structures to compute similarity of molecule structures. *CEJOR Cent*. Eur. J. Oper. Res. 22, 457–473 (2014).

31. Zhang, C., Shine, M., Pyle, A. M. & Zhang, Y. US-align: universal structure alignments of proteins, nucleic acids, and macromolecular complexes. Nat. Methods 19, 1109–1115 (2022).

32. Zemla, A. LGA: A method for finding 3D similarities in protein structures. Nucleic Acids Res. 31, 3370–3374 (2003).

33. Magnus, M., Antczak, M., Zok, T., Wiedemann, J., Lukasiak, P., Cao, Y., Bujnicki, J. M., Westhof, E., Szachniuk, M. & Miao, Z. RNA-Puzzles toolkit: a computational resource of RNA 3D structure benchmark datasets, structure manipulation, and evaluation tools. Nucleic Acids Res. 48, 576–588 (2020).

34. Waleń, T., Chojnowski, G., Gierski, P. & Bujnicki, J. M. ClaRNA: a classifier of contacts in RNA 3D structures based on a comparative analysis of various classification schemes. Nucleic Acids Res. 42, e151 (2014).

35. Mariani, V., Biasini, M., Barbato, A. & Schwede, T. lDDT: a local superposition-free score for comparing protein structures and models using distance difference tests. Bioinformatics 29, 2722–2728 (2013).

36. Biasini, M., Mariani, V., Haas, J., Scheuber, S., Schenk, A. D., Schwede, T. & Philippsen, A. OpenStructure: a flexible software framework for computational structural biology. Bioinformatics 26, 2626–2628 (2010).

37. Liebschner, D., Afonine, P. V., Baker, M. L., Bunkóczi, G., Chen, V. B., Croll, T. I., Hintze, B., Hung, L. W., Jain, S., McCoy, A. J., Moriarty, N. W., Oeffner, R. D., Poon, B. K., Prisant, M. G., Read, R. J., Richardson, J. S., Richardson, D. C., Sammito, M. D., Sobolev, O. V., Stockwell, D. H., Terwilliger, T. C., Urzhumtsev, A. G., Videau, L. L., Williams, C. J. & Adams, P. D. Macromolecular structure determination using X-rays, neutrons and electrons: recent developments in Phenix. Acta Crystallogr D Struct Biol 75, 861–877 (2019).

38. Kwon, S., Won, J., Kryshtafovych, A. & Seok, C. Assessment of protein model structure accuracy estimation in CASP14: Old and new challenges. Proteins 89, 1940–1948 (2021).

39. Beton, J. G., Cragnolini, T., Kaleel, M., Mulvaney, T., Sweeney, A. & Topf, M. Integrating model simulation tools and cryoLelectron microscopy. Wiley Interdiscip. Rev. Comput. Mol. Sci. 13, e1642 (2023).

40. Watkins, A. M., Rangan, R. & Das, R. Using Rosetta for RNA homology modeling. Methods Enzymol. 623, 177–207 (2019).

41. Pettersen, E. F., Goddard, T. D., Huang, C. C., Meng, E. C., Couch, G. S., Croll, T. I., Morris, J. H. & Ferrin, T. E. UCSF ChimeraX: Structure visualization for researchers, educators, and developers. Protein Sci. 30, 70–82 (2021).

42. Liebschner, D., Afonine, P. V., Baker, M. L., Bunkóczi, G., Chen, V. B., Croll, T. I., Hintze, B., Hung, L. W., Jain, S., McCoy, A. J., Moriarty, N. W., Oeffner, R. D., Poon, B. K., Prisant, M. G., Read, R. J., Richardson, J. S., Richardson, D. C., Sammito, M. D., Sobolev, O. V., Stockwell, D. H., Terwilliger, T. C., Urzhumtsev, A. G., Videau, L. L., Williams, C. J. & Adams, P. D. Macromolecular structure determination using X-rays, neutrons and electrons: recent developments in Phenix. Acta Crystallogr D Struct Biol 75, 861–877 (2019).

43. Cragnolini, T., Sahota, H., Joseph, A. P., Sweeney, A., Malhotra, S., Vasishtan, D. & Topf, M. TEMPy2: a Python library with improved 3D electron microscopy density-fitting and validation workflows. Acta Crystallogr D Struct Biol 77, 41–47 (2021).

44. Lagerstedt, I., Moore, W. J., Patwardhan, A., Sanz-García, E., Best, C., Swedlow, J. R. & Kleywegt, G. J. Web-based visualisation and analysis of 3D electron-microscopy data from EMDB and PDB. J. Struct. Biol. 184, 173–181 (2013).

45. Pintilie, G., Zhang, K., Su, Z., Li, S., Schmid, M. F. & Chiu, W. Measurement of atom resolvability in cryo-EM maps with Q-scores. Nat. Methods 17, 328–334 (2020).

46. McCoy, A. J., Grosse-Kunstleve, R. W., Adams, P. D., Winn, M. D., Storoni, L. C. & Read, R. J. Phaser crystallographic software. J. Appl. Crystallogr. 40, 658–674 (2007).

47. Winn, M. D., Ballard, C. C., Cowtan, K. D., Dodson, E. J., Emsley, P., Evans, P. R., Keegan, R. M., Krissinel, E. B., Leslie, A. G. W., McCoy, A., McNicholas, S. J., Murshudov, G. N., Pannu, N. S., Potterton, E. A., Powell, H. R., Read, R. J., Vagin, A. & Wilson, K. S. Overview of the CCP4 suite and current developments. Acta Crystallogr. D Biol. Crystallogr. 67, 235–242 (2011).

48. Krissinel, E., Lebedev, A. A., Uski, V., Ballard, C. B., Keegan, R. M., Kovalevskiy, O., Nicholls, R. A., Pannu, N. S., Skubák, P., Berrisford, J., Fando, M., Lohkamp, B., Wojdyr, M., Simpkin, A. J., Thomas, J. M. H., Oliver, C., Vonrhein, C., Chojnowski, G., Basle, A., Purkiss, A., Isupov, M. N., McNicholas, S., Lowe, E., Triviño, J., Cowtan, K., Agirre, J., Rigden, D. J., Uson, I., Lamzin, V., Tews, I., Bricogne, G., Leslie, A. G. W. & Brown, D. G. CCP4 Cloud for structure determination and project management in macromolecular crystallography. Acta Crystallogr D Struct Biol 78, 1079–1089 (2022).

49. Vagin, A. & Teplyakov, A. Molecular replacement with MOLREP. Acta Crystallogr. D Biol. Crystallogr. 66, 22–25 (2010).

50. Theobald, D. L. & Wuttke, D. S. THESEUS: maximum likelihood superpositioning and analysis of macromolecular structures. Bioinformatics 22, 2171–2172 (2006).

51. Jumper, J., Evans, R., Pritzel, A., Green, T., Figurnov, M., Ronneberger, O., Tunyasuvunakool, K., Bates, R., Žídek, A., Potapenko, A., Bridgland, A., Meyer, C., Kohl, S. A. A., Ballard, A. J., Cowie, A., Romera-Paredes, B., Nikolov, S., Jain, R., Adler, J., Back, T., Petersen, S., Reiman, D., Clancy, E., Zielinski, M., Steinegger, M., Pacholska, M., Berghammer, T., Bodenstein, S., Silver, D., Vinyals, O., Senior, A. W., Kavukcuoglu, K., Kohli, P. & Hassabis, D. Highly accurate protein structure prediction with AlphaFold. Nature 596, 583–589 (2021).

52. Mirdita, M., Schütze, K., Moriwaki, Y., Heo, L., Ovchinnikov, S. & Steinegger, M. ColabFold: making protein folding accessible to all. Nat. Methods 19, 679–682 (2022).

53. Simpkin, A. J., Elliott, L. G., Stevenson, K., Krissinel, E., Rigden, D. & Keegan, R. M. Slice’N’Dice: Maximising the value of predicted models for structural biologists. bioRxiv (2022). doi:10.1101/2022.06.30.497974

54. Pedregosa, F., Varoquaux, G., Gramfort, A., Michel, V., Thirion, B., Grisel, O., Blondel, M., Müller, A., Nothman, J., Louppe, G., Prettenhofer, P., Weiss, R., Dubourg, V., Vanderplas, J., Passos, A., Cournapeau, D., Brucher, M., Perrot, M. & Duchesnay, É. Scikit-learn: Machine Learning in Python. (2012). doi:10.48550/ARXIV.1201.0490

55. Read, R. J. & Chavali, G. Assessment of CASP7 predictions in the high accuracy template-based modeling category. Proteins 69 **Suppl 8**, 27–37 (2007).

56. Kinch, L. N., Pei, J., Kryshtafovych, A., Schaeffer, R. D. & Grishin, N. V. Topology evaluation of models for difficult targets in the 14th round of the critical assessment of protein structure prediction (CASP14). Proteins 89, 1673–1686 (2021).

57. Haas, J., Barbato, A., Behringer, D., Studer, G., Roth, S., Bertoni, M., Mostaguir, K., Gumienny, R. & Schwede, T. Continuous Automated Model EvaluatiOn (CAMEO) complementing the critical assessment of structure prediction in CASP12. Proteins 86, 387–398 (2018).

58. Robin, X., Haas, J., Gumienny, R., Smolinski, A., Tauriello, G. & Schwede, T. Continuous Automated Model EvaluatiOn (CAMEO)-Perspectives on the future of fully automated evaluation of structure prediction methods. Proteins 89, 1977–1986 (2021).

59. Przytula-Mally, A. I., Engilberge, S., Johannsen, S., Olieric, V., Masquida, B. & Sigel, R. K. O. Anticodon-like loop-mediated dimerization in the crystal structures of HdV-like CPEB3 ribozymes. bioRxiv 2022.09.22.508989 (2022). doi:10.1101/2022.09.22.508989

60. Vallina, N. S., McRae, E. K. S., Hansen, B. K., Boussebayle, A. & Andersen, E. S. RNA origami scaffolds as a cryo-EM tool for investigating aptamer-ligand binding of a Broccoli-Pepper FRET pair. bioRxiv 2022.08.25.505116 (2022). doi:10.1101/2022.08.25.505116

61. McRae, E. K. S., Rasmussen, H. Ø., Liu, J., Bøggild, A., Nguyen, M. T. A., Sampedro Vallina, N., Boesen, T., Pedersen, J. S., Ren, G., Geary, C. & Others. Structure, folding and flexibility of co-transcriptional RNA origami. Nat. Nanotechnol. 1–10 (2023).

62. Chen, J.-H., Yajima, R., Chadalavada, D. M., Chase, E., Bevilacqua, P. C. & Golden, B. L. A 1.9 A crystal structure of the HDV ribozyme precleavage suggests both Lewis acid and general acid mechanisms contribute to phosphodiester cleavage. Biochemistry 49, 6508– 6518 (2010).

63. Jenkins, J. L., Krucinska, J., McCarty, R. M., Bandarian, V. & Wedekind, J. E. Comparison of a preQ1 riboswitch aptamer in metabolite-bound and free states with implications for gene regulation. J. Biol. Chem. 286, 24626–24637 (2011).

64. Chen, K., Zhou, Y., Wang, S. & Xiong, P. RNA tertiary structure modeling with BRiQ potential in CASP15. bioRxiv 2023.05.26.542548 (2023). doi:10.1101/2023.05.26.542548

65. Cragnolini, T., Kryshtafovych, A. & Topf, M. Cryo-EM targets in CASP14. Proteins 89, 1949–1958 (2021).

66. Vasishtan, D. & Topf, M. Scoring functions for cryoEM density fitting. J. Struct. Biol. 174, 333–343 (2011).

67. Joseph, A. P., Lagerstedt, I., Patwardhan, A., Topf, M. & Winn, M. Improved metrics for comparing structures of macromolecular assemblies determined by 3D electron-microscopy. J. Struct. Biol. 199, 12–26 (2017).

68. Vallina, N. S., McRae, E. K. S., Geary, C. & Andersen, E. S. An RNA Paranemic Crossover Triangle as A 3D Module for Cotranscriptional Nanoassembly. Small 2204651 Preprint at 10.1002/smll.202204651 (2022)

69. Bonilla, S. L., Vicens, Q. & Kieft, J. S. Cryo-EM reveals an entangled kinetic trap in the folding of a catalytic RNA. Sci Adv 8, eabq4144 (2022).

70. Li, S., Palo, M. Z., Pintilie, G., Zhang, X., Su, Z., Kappel, K., Chiu, W., Zhang, K. & Das, R. Topological crossing in the misfolded Tetrahymena ribozyme resolved by cryo-EM. Proc. Natl. Acad. Sci. U. S. A. 119, e2209146119 (2022).

71. Huang, L., Wang, J., Watkins, A. M., Das, R. & Lilley, D. M. J. Structure and ligand binding of the glutamine-II riboswitch. Nucleic Acids Res. 47, 7666–7675 (2019).

72. Murshudov, G. N., Skubák, P., Lebedev, A. A., Pannu, N. S., Steiner, R. A., Nicholls, R. A., Winn, Long, F. & Vagin, A. A. REFMAC5 for the refinement of macromolecular crystal structures. Acta Crystallogr. D Biol. Crystallogr. 67, 355–367 (2011).

73. Theobald, D. L. & Wuttke, D. S. Accurate Structural Correlations from Maximum Likelihood Superpositions. PLoS Comput. Biol. 4, e43 (2008).

74. Bibby, J., Keegan, R. M., Mayans, O., Winn, M. D. & Rigden, D. J. AMPLE: a cluster-and-truncate approach to solve the crystal structures of small proteins using rapidly computed ab initio models. Acta Crystallogr. D Biol. Crystallogr. 68, 1622–1631 (2012).

75. Oeffner, R. D., Afonine, P. V., Millán, C., Sammito, M., Usón, I., Read, R. J. & McCoy, A. J. On the application of the expected log-likelihood gain to decision making in molecular replacement. Acta Crystallogr D Struct Biol 74, 245–255 (2018).

76. Shen, T., Hu, Z., Peng, Z., Chen, J., Xiong, P., Hong, L., Zheng, L., Wang, Y., King, I., Wang, S., Sun, S. & Li, Y. E2Efold-3D: End-to-End Deep Learning Method for accurate de novo RNA 3D Structure Prediction. (2022). at <http://arxiv.org/abs/2207.01586>

77. Pearce, R., Omenn, G. S. & Zhang, Y. De Novo RNA Tertiary Structure Prediction at Atomic Resolution Using Geometric Potentials from Deep Learning. bioRxiv 2022.05.15.491755 (2022). doi:10.1101/2022.05.15.491755

78. Strobel, E. J., Yu, A. M. & Lucks, J. B. High-throughput determination of RNA structures. Nat. Rev. Genet. 19, 615–634 (2018).

79. Cossio, P., Rohr, D., Baruffa, F., Rampp, M., Lindenstruth, V. & Hummer, G. BioEM: GPU-accelerated computing of Bayesian inference of electron microscopy images. Comput. Phys. Commun. 210, 163–171 (2017).

80. Kryshtafovych, A., Antczak, M., Szachniuk, M., Zok, T., Kretsch, R. C., Rangan, R., Pham, P., Das, R., Robin, X., Studer, G., Durairaj, J., Eberhardt, J., Sweeney, A., Topf, M., Schwede, T., Fidelis, K. & Moult, J. New prediction categories in CASP15. Proteins (2023). doi:10.1002/prot.26515

81. Leontis, N. B. & Westhof, E. Geometric nomenclature and classification of RNA base pairs. RNA 7, 499–512 (2001).

