## Supplemental Information for "Assessment of three-dimensional RNA structure prediction in CASP15"


Rhiju Das[1,2,3,*,‡] https://orcid.org/0000-0001-7497-0972
Rachael C. Kretsch[2,*] https://orcid.org/0000-0002-6935-518X
Adam Simpkin[4,†] https://orcid.org/0000-0003-1883-9376
Thomas Mulvaney[5,6,†] https://orcid.org/0000-0002-4373-6160
Phillip Pham[1] https://orcid.org/0000-0002-0240-6384
Ramya Rangan[2] https://orcid.org/0000-0002-0960-0825
Fan Bu[7,8] https://orcid.org/0009-0000-1903-2270
Ronan Keegan[4,9] https://orcid.org/0000-0002-9495-0431
Maya Topf[5,6] https://orcid.org/0000-0002-8185-1215
Daniel Rigden[4] https://orcid.org/0000-0002-7565-8937
Zhichao Miao[10,11,‡] https://orcid.org/0000-0002-5777-9815
Eric Westhof[12,‡] https://orcid.org/0000-0002-6172-5422

[1]Department of Biochemistry and [2]Biophysics Program, Stanford University School of Medicine, CA USA, [3]Howard Hughes Medical Institute, Stanford University, CA USA, [4]Institute of Systems, Molecular & Integrative Biology, The University of Liverpool, UK, [5]Centre for Structural Systems Biology (CSSB), Leibniz-Institut für Virologie (LIV) and [6]University Medical Center Hamburg-Eppendorf (UKE), Hamburg, Germany, [7]Guangzhou Laboratory, Guangzhou International Bio Island, Guangzhou 510005, China, [8]Division of Life Sciences and Medicine,University of Science and Technology of China, Hefei 230036, Anhui, China, [9]Life Science, Diamond Light Source, Harwell Science, UK, [10]GMU-GIBH Joint School of Life Sciences and [11]The Guangdong-Hong Kong-Macau Joint Laboratory for Cell Fate Regulation and Diseases, Guangzhou Laboratory, Guangzhou Medical University, and [12]Architecture et Réactivité de l'ARN, Institut de Biologie Moléculaire et Cellulaire du CNRS, Université de Strasbourg, F-67084, Strasbourg, France. [*]Equally contributing authors. [†]Equally contributing authors. [‡]Correspondence to:, and.


This Supplemental Information file contains one Supplemental Table and six Supplemental Figures.

| Model | Best RMSD[a] | | | | Model 1 RMSD[b] | | | | Best DI[c] | | | | Model 1 DI[c] | | | | Best INF[d] | | | | Model 1 INF[d] | | | |
|---|---|---|---|---|---|---|---|---|---|---|---|---|---|---|---|---|---|---|---|---|---|---|---|---|
| Rank | 1st | 2nd | 3rd | sum[e] | 1st | 2nd | 3rd | sum[e] | 1st | 2nd | 3rd | sum[e] | 1st | 2nd | 3rd | sum[e] | 1st | 2nd | 3rd | sum[e] | 1st | 2nd | 3rd | sum[e] |
| TS232 | 6 | 2 | 0 | 22 | 5 | 0 | 2 | 17 | 6 | 2 | 0 | 22 | 6 | 0 | 1 | 19 | 6 | 1 | 1 | 21 | 3 | 3 | 0 | 15 |
| TS287 | 1 | 2 | 2 | 9 | 1 | 3 | 2 | 11 | 1 | 3 | 2 | 11 | 2 | 5 | 0 | 16 | 2 | 0 | 3 | 9 | 4 | 2 | 0 | 16 |
| TS128 | 1 | 1 | 2 | 7 | 2 | 1 | 0 | 8 | 1 | 0 | 1 | 4 | 1 | 1 | 1 | 6 | 0 | 0 | 0 | 0 | 0 | 0 | 0 | 0 |
| TS081 | 0 | 1 | 1 | 3 | 2 | 1 | 1 | 9 | 0 | 1 | 3 | 5 | 1 | 3 | 1 | 10 | 2 | 2 | 2 | 12 | 2 | 1 | 2 | 10 |
| TS229 | 2 | 0 | 1 | 7 | 0 | 0 | 0 | 0 | 2 | 0 | 0 | 6 | 0 | 0 | 0 | 0 | 0 | 0 | 0 | 0 | 0 | 0 | 0 | 0 |
| TS416 | 0 | 2 | 2 | 6 | 0 | 1 | 0 | 2 | 0 | 1 | 1 | 3 | 0 | 1 | 0 | 2 | 0 | 0 | 0 | 0 | 0 | 1 | 0 | 2 |
| TS239 | 2 | 0 | 0 | 6 | 0 | 0 | 0 | 0 | 2 | 0 | 0 | 6 | 0 | 0 | 0 | 0 | 0 | 0 | 0 | 0 | 0 | 0 | 0 | 0 |
| TS439 | 2 | 0 | 0 | 6 | 0 | 0 | 0 | 0 | 2 | 0 | 0 | 6 | 0 | 0 | 0 | 0 | 0 | 0 | 0 | 0 | 0 | 0 | 0 | 0 |
| TS110 | 1 | 0 | 0 | 3 | 0 | 0 | 0 | 0 | 1 | 0 | 0 | 3 | 0 | 0 | 1 | 1 | 0 | 0 | 0 | 0 | 0 | 0 | 0 | 0 |
| TS285 | 1 | 0 | 0 | 3 | 1 | 0 | 0 | 3 | 1 | 0 | 0 | 3 | 1 | 0 | 0 | 3 | 0 | 0 | 0 | 0 | 1 | 0 | 0 | 3 |
| TS456 | 0 | 1 | 0 | 2 | 0 | 1 | 0 | 2 | 0 | 1 | 0 | 2 | 0 | 1 | 0 | 2 | 0 | 0 | 1 | 1 | 0 | 0 | 0 | 0 |
| TS054 | 0 | 1 | 0 | 2 | 0 | 1 | 0 | 2 | 0 | 1 | 0 | 2 | 1 | 0 | 0 | 3 | 0 | 1 | 1 | 3 | 0 | 0 | 2 | 2 |
| TS347 | 0 | 1 | 0 | 2 | 0 | 1 | 0 | 2 | 0 | 1 | 0 | 2 | 0 | 1 | 0 | 2 | 0 | 0 | 1 | 1 | 0 | 0 | 0 | 0 |
| TS489 | 0 | 0 | 0 | 0 | 1 | 0 | 1 | 4 | 0 | 0 | 0 | 0 | 0 | 0 | 1 | 1 | 0 | 0 | 0 | 0 | 0 | 0 | 0 | 0 |
| TS470 | 0 | 0 | 0 | 0 | 1 | 0 | 1 | 4 | 0 | 0 | 0 | 0 | 0 | 0 | 1 | 1 | 0 | 0 | 0 | 0 | 0 | 0 | 0 | 0 |
| TS227 | 0 | 0 | 0 | 0 | 0 | 1 | 0 | 2 | 0 | 0 | 0 | 0 | 0 | 0 | 0 | 0 | 0 | 0 | 0 | 0 | 0 | 0 | 0 | 0 |
| TS147 | 0 | 0 | 0 | 0 | 0 | 1 | 0 | 2 | 0 | 0 | 0 | 0 | 0 | 0 | 0 | 0 | 0 | 0 | 0 | 0 | 0 | 0 | 0 | 0 |
| TS119 | 0 | 0 | 0 | 0 | 0 | 1 | 0 | 2 | 0 | 0 | 0 | 0 | 0 | 1 | 1 | 3 | 0 | 1 | 1 | 3 | 1 | 0 | 0 | 3 |
| TS325 | 0 | 0 | 0 | 0 | 0 | 1 | 0 | 2 | 0 | 1 | 0 | 2 | 0 | 1 | 0 | 2 | 0 | 0 | 1 | 1 | 0 | 0 | 0 | 0 |
| TS392 | 0 | 0 | 0 | 0 | 0 | 1 | 0 | 2 | 0 | 0 | 0 | 0 | 0 | 0 | 0 | 0 | 0 | 0 | 0 | 0 | 0 | 1 | 0 | 2 |
| TS434 | 0 | 0 | 0 | 0 | 0 | 1 | 0 | 2 | 0 | 0 | 0 | 0 | 0 | 0 | 0 | 0 | 1 | 1 | 0 | 5 | 0 | 0 | 1 | 1 |
| TS035 | 0 | 0 | 0 | 0 | 0 | 0 | 0 | 0 | 0 | 1 | 0 | 2 | 0 | 0 | 0 | 0 | 0 | 1 | 1 | 3 | 0 | 0 | 2 | 2 |
| TSR01 | 0 | 0 | 0 | 0 | 0 | 0 | 0 | 0 | 0 | 0 | 2 | 2 | 0 | 0 | 1 | 1 | 0 | 0 | 1 | 1 | 0 | 1 | 0 | 2 |
| TS125 | 0 | 0 | 0 | 0 | 0 | 0 | 0 | 0 | 0 | 0 | 1 | 1 | 0 | 0 | 1 | 1 | 0 | 1 | 0 | 2 | 0 | 0 | 1 | 1 |
| TS235 | 0 | 0 | 0 | 0 | 0 | 0 | 0 | 0 | 0 | 0 | 0 | 0 | 0 | 0 | 1 | 1 | 0 | 0 | 0 | 0 | 0 | 0 | 1 | 1 |
| TS076 | 0 | 0 | 0 | 0 | 0 | 0 | 0 | 0 | 0 | 0 | 0 | 0 | 0 | 0 | 1 | 1 | 0 | 0 | 0 | 0 | 0 | 0 | 0 | 0 |
| TS444 | 0 | 0 | 0 | 0 | 0 | 0 | 0 | 0 | 0 | 0 | 0 | 0 | 0 | 0 | 1 | 1 | 0 | 2 | 0 | 4 | 0 | 1 | 1 | 3 |
| TS248 | 0 | 0 | 0 | 0 | 0 | 0 | 0 | 0 | 0 | 0 | 0 | 0 | 0 | 0 | 0 | 0 | 0 | 0 | 1 | 1 | 0 | 0 | 1 | 1 |
| TS185 | 0 | 0 | 0 | 0 | 0 | 0 | 0 | 0 | 0 | 0 | 0 | 0 | 0 | 0 | 0 | 0 | 0 | 0 | 0 | 0 | 1 | 0 | 0 | 3 |

[a] Best RMSD reached for all the models submitted.

[b] RMSD reached by the model ranked #1 by the group.

[c] Same as [a,b] for Deformation Index (DI).

[d] Same as [a,b] for Interaction Network Fidelity (INF).

[e] For each group, the number of times the submitted models were ranked 1st, 2nd, or 3rd were counted and the weighted sum indicated in SUM (with a weight of 3 for 1st, 2 for 2nd and 1 for 3rd ranks).

**Supplementary Table 1.** Scores reached by the modeling groups using the different RNA-Puzzles metrics and assessment processes as indicated in the column Model.

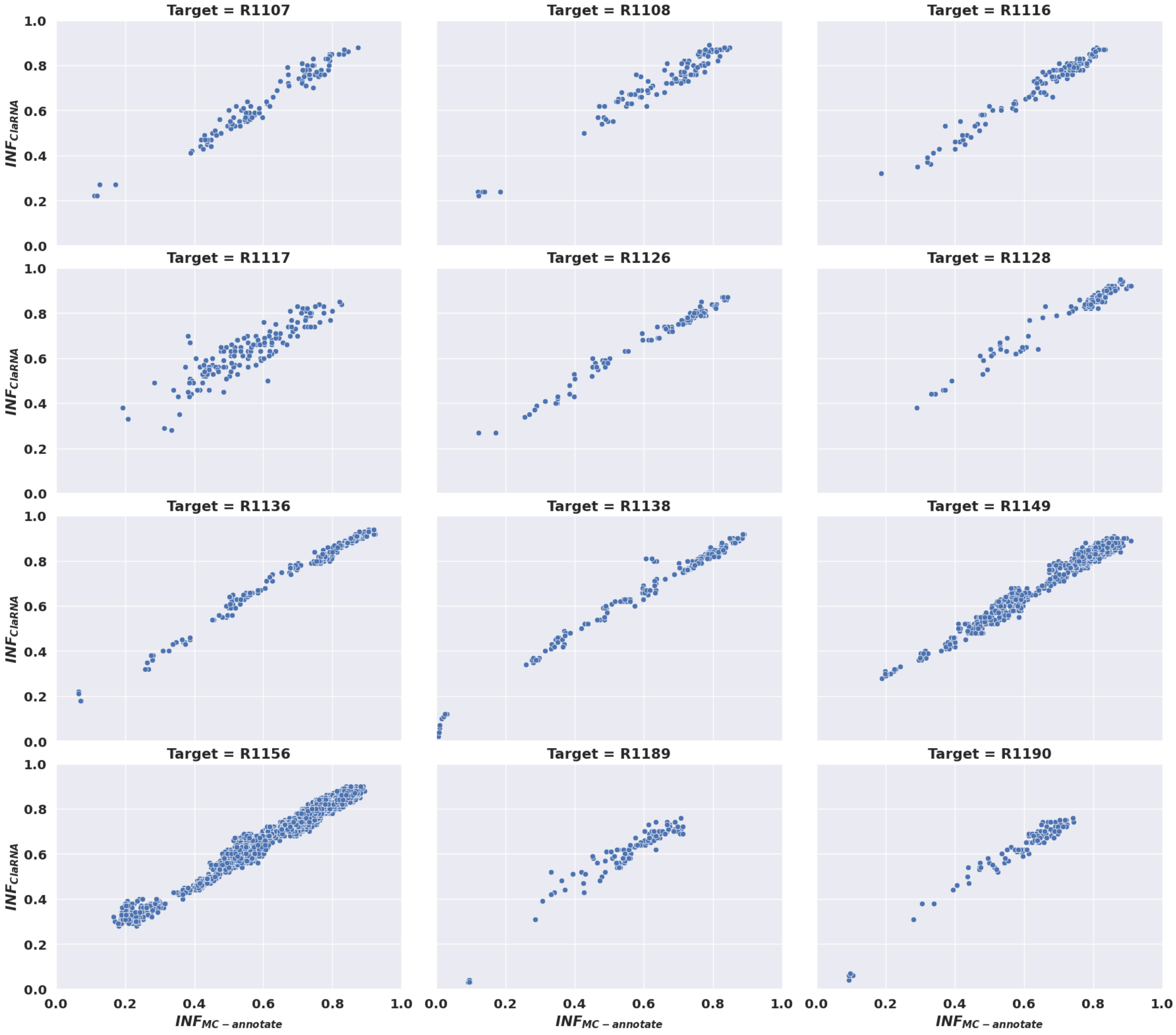


**Supplemental Figure 1. Comparison of two tools to calculate interaction network fidelity (INF) of RNA.** Comparison of INF computed using MC-annotate vs. using ClaRNA.

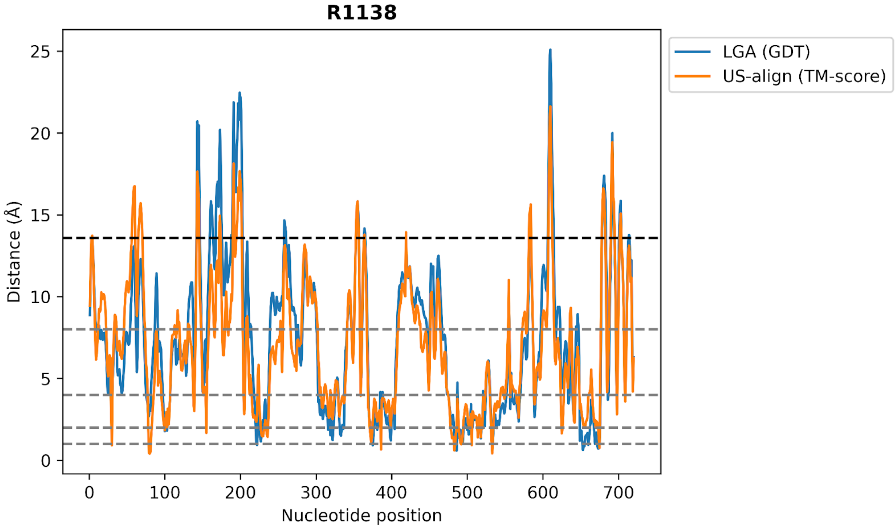


**Supplemental Figure 2. How a good prediction can get high TM-score but low GDT_TS.** For the R1138 model 4 submitted by AIchemy_RNA2 (R1138TS232_4) and the cryo-EM R1138 structure, the residue-residue distances between the C3' and C4' atoms were calculated using the superimposed coordinates determined by US-align and LGA, respectively; the traces are similar. The black dashed line represents the (soft) distance threshold used in US-align to compute TM-score ($d_0$= 13.59 Å), which is set based on the molecule length; for this 720-nucleotide target the $d_0$ value is large and most residues align within the threshold, leading to a high TM-score for this target. In contrast, the gray lines indicate the threshold values used in GDT_TS (1 Å, 2 Å, 4 Å, and 8 Å). These threshold values do not change with molecule length and so do not take into account the increased flexibility expected for longer RNA molecules, leading to small GDT_TS values for this target.

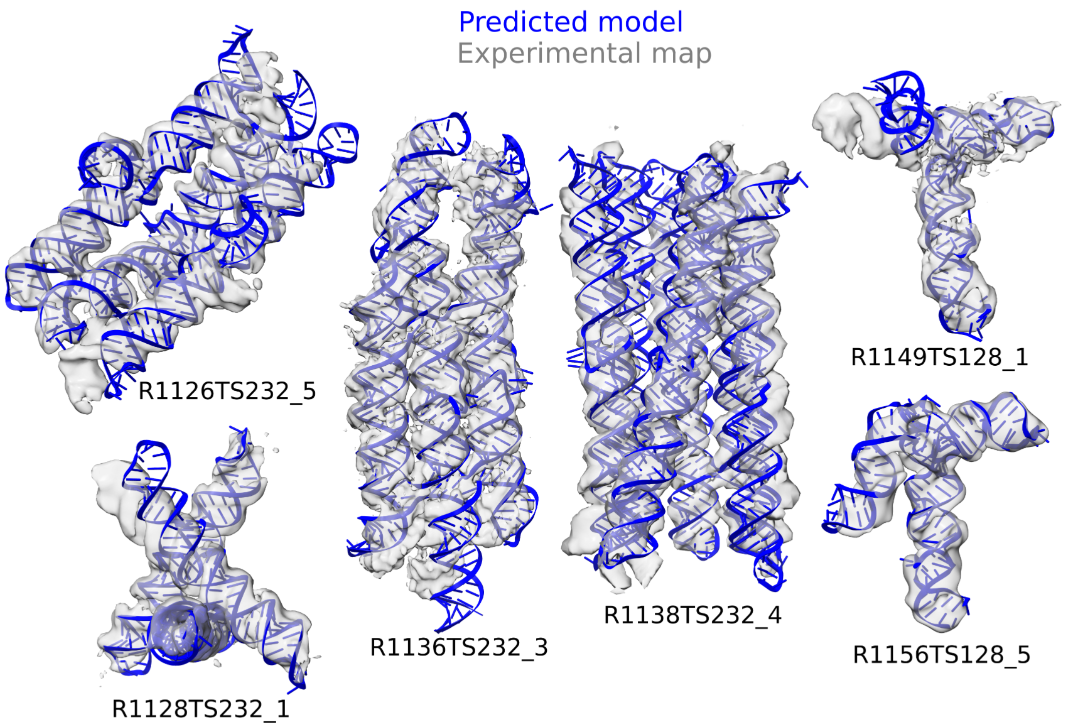


**Supplemental Figure 3. Fits of CASP15 RNA models to EM maps.** The best fitting (by $Z_{EM}$) predicted model (blue) fit into an experimental cryo-EM map for each target (gray).

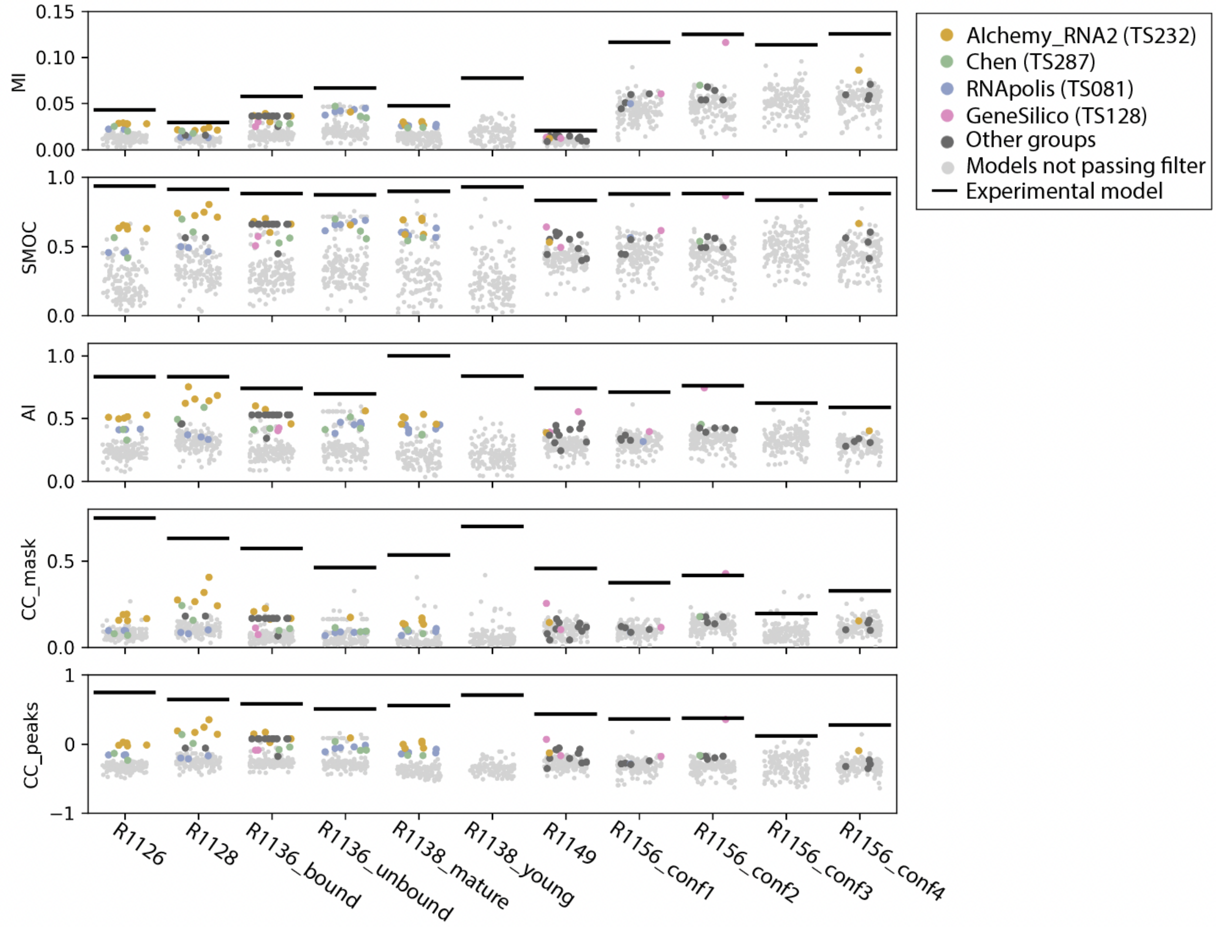


**Supplemental Figure 4. EM metrics for all targets.** Scores for all models submitted for all targets are depicted. Models passing the RMSD filter are indicated with larger dots, colored if submitted by top performing groups and dark gray otherwise. The black line indicates the EM metric scores for the experimentally determined model.

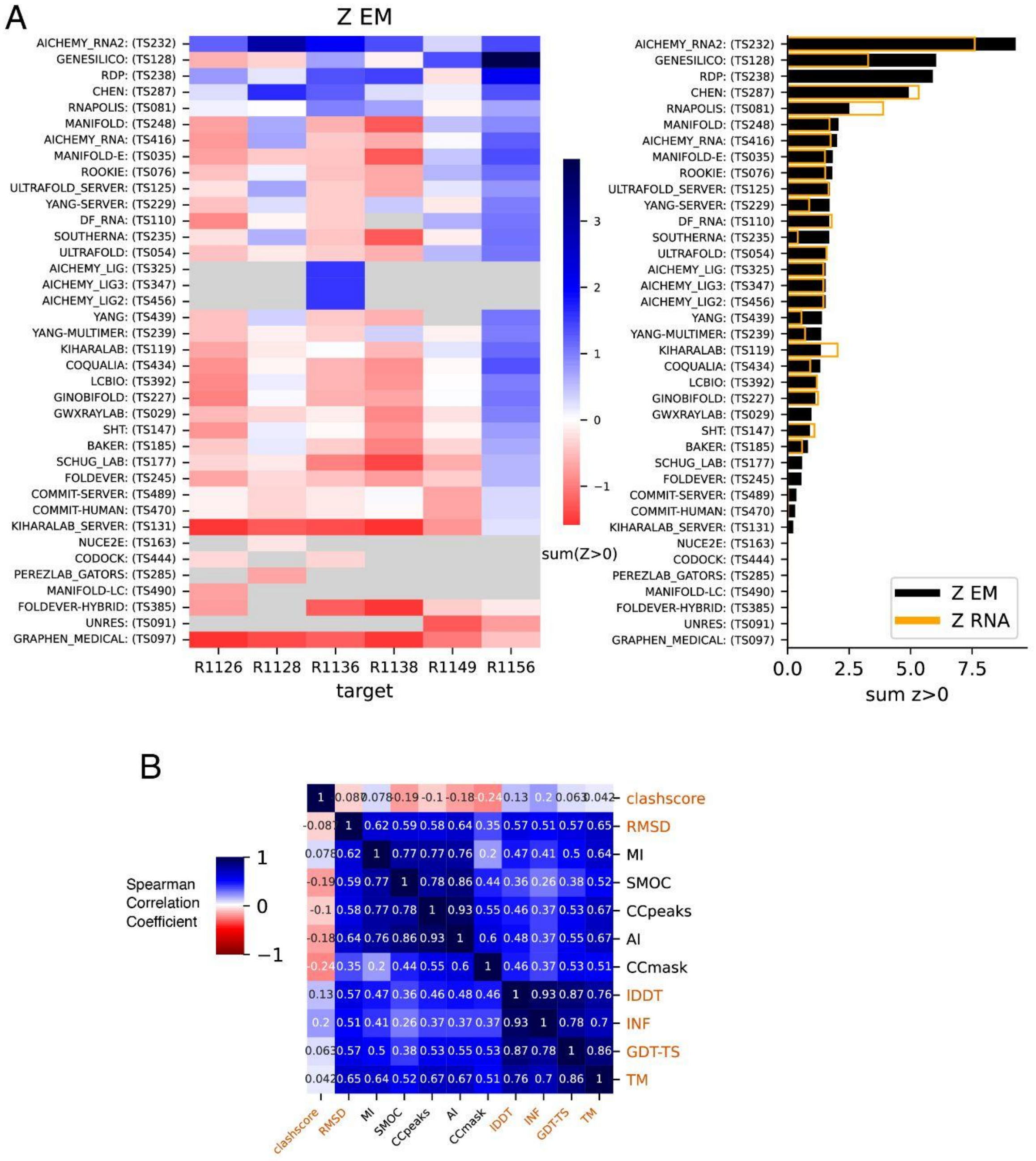


**Supplemental Figure 5. Map-to-model analysis over all submitted models without RMSD filtering**. (A) Z-scores for all models for RNA cryo-EM targets and ranking according to $Z_{EM}$ (black), in orange are the $Z_{RNA}$ scores for comparison. (B) the Spearman correlation between metrics used in $Z_{RNA}$, $Z_{EM}$, as well as RMSD for all models from cryo-EM targets. RMSD and clashscore were multiplied by -1 before calculating the correlation so that higher scores corresponded to better accuracy for all metrics.

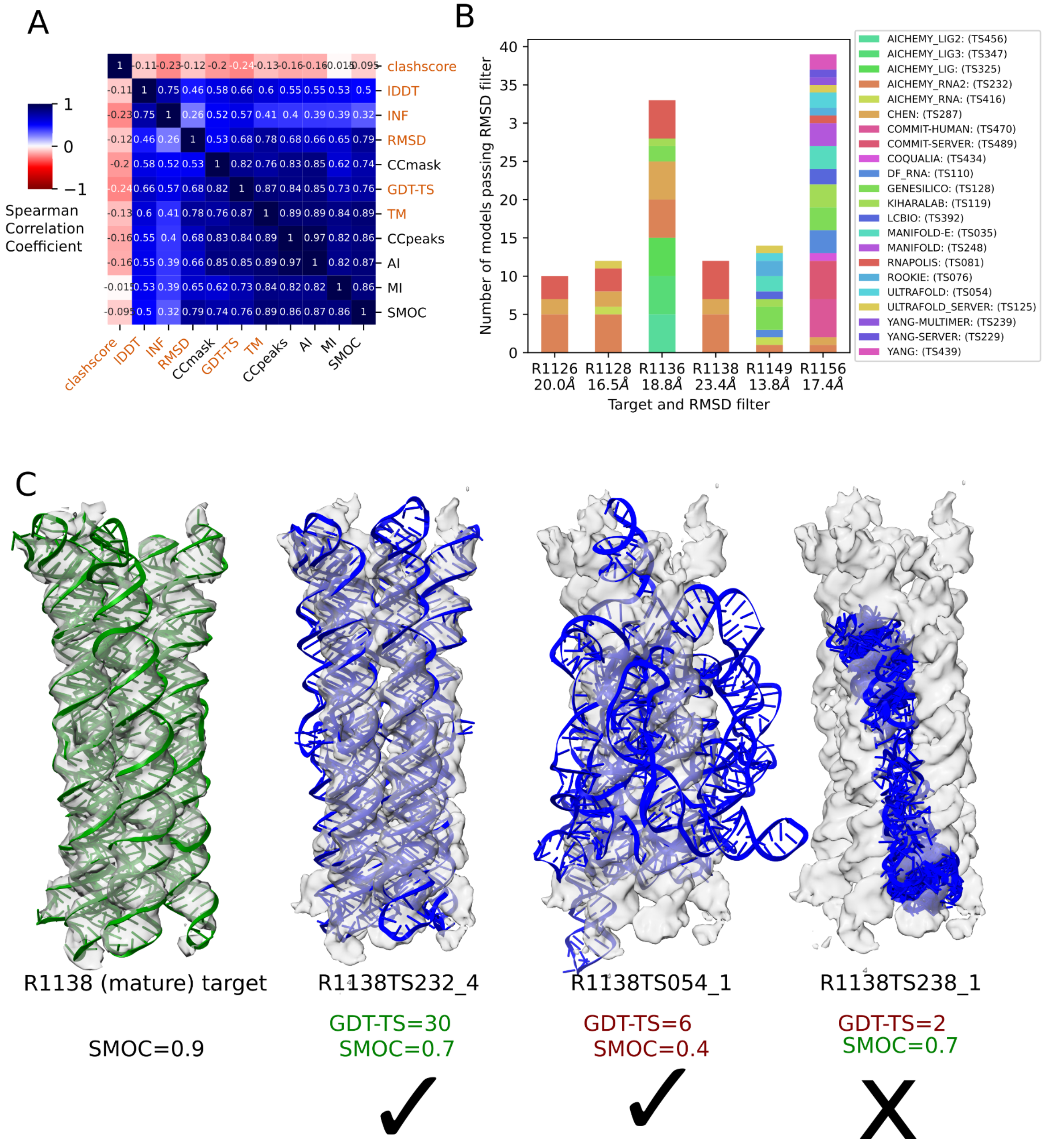


**Supplemental Figure 6. Map-to-model analysis and RMSD filtering.** (A) the Spearman correlation between metrics used in $Z_{RNA}$, $Z_{EM}$, as well as RMSD, computed for all models passing the RMSD filter. RMSD and clashscore were multiplied by -1 before calculating the correlation so that higher scores corresponded to better accuracy for all metrics. (B) The number of models which have RMSD to target less than the filter cutoff; these models were used in the final EM ranking. (C) An example of when EM metrics can be misleading. Reference structure in green, experimental map in grey, predicted models in blue. GDT-TS is reported as an example of model-to-model metric and SMOC as an example of map-to-model metrics. Green scores are seen as “good” while red are “poor” scores

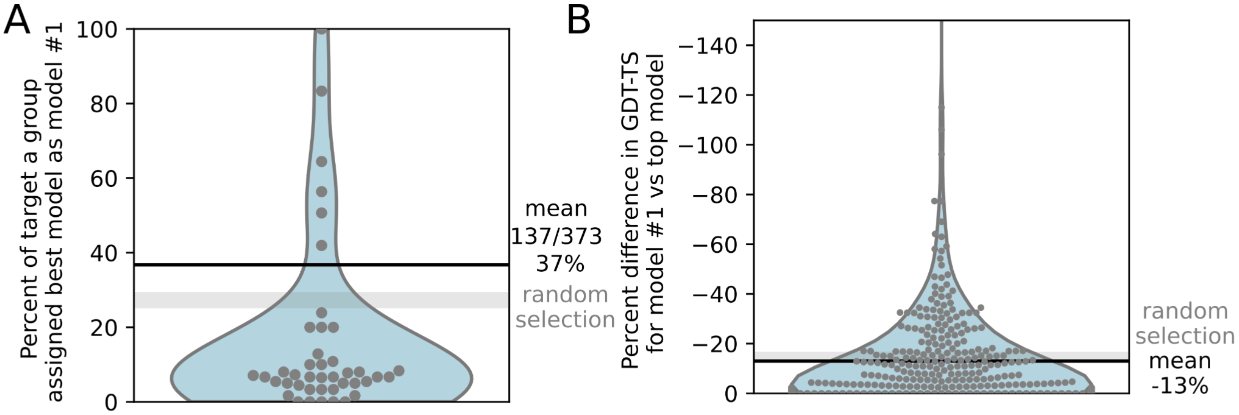


**Supplemental Figure 7. Comparison of groups' top model by GDT-TS and the model they selected as model #1**. (A) For each group, the percent of targets they participated in where their best model by GDT-TS (out of up to five models submitted) was assigned as model #1. (B) For all targets and all groups, the percent difference in GDT-TS from model #1 to the top model for that group. The mean values over CASP groups in (A,B) are shown as black lines. For comparison, the gray bars in (A,B) mark the 95% confidence interval for values from random shuffling to select "model #1" (1,000 and 10,000 bootstraps respectively).

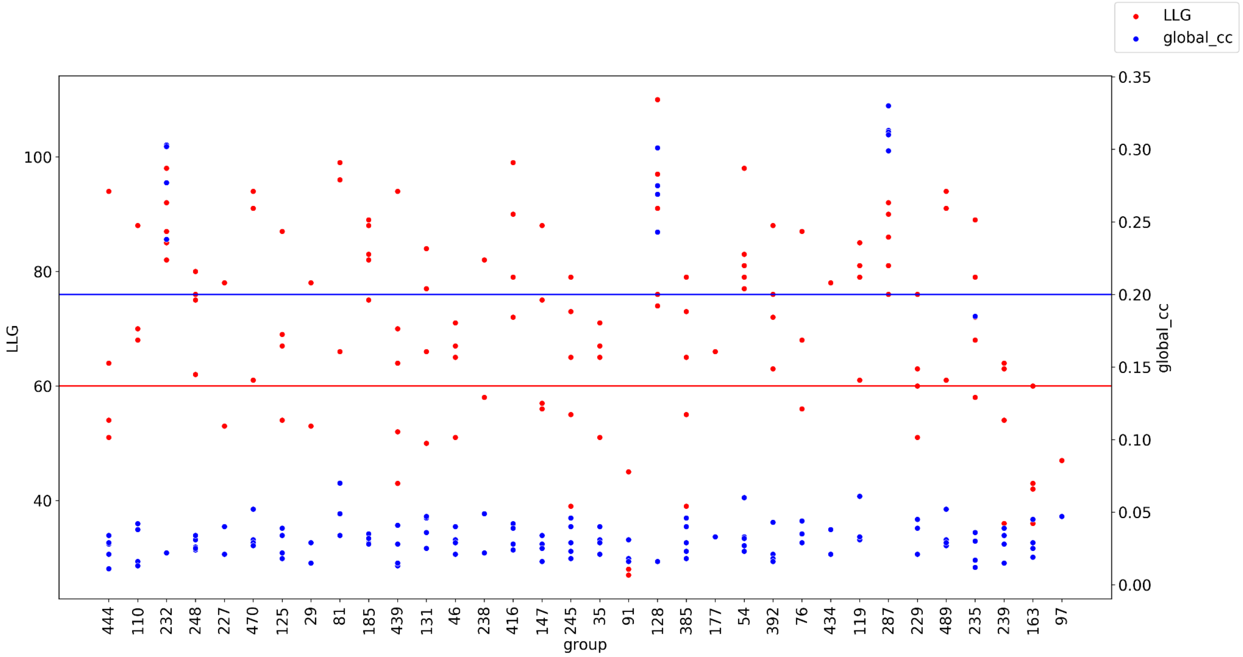


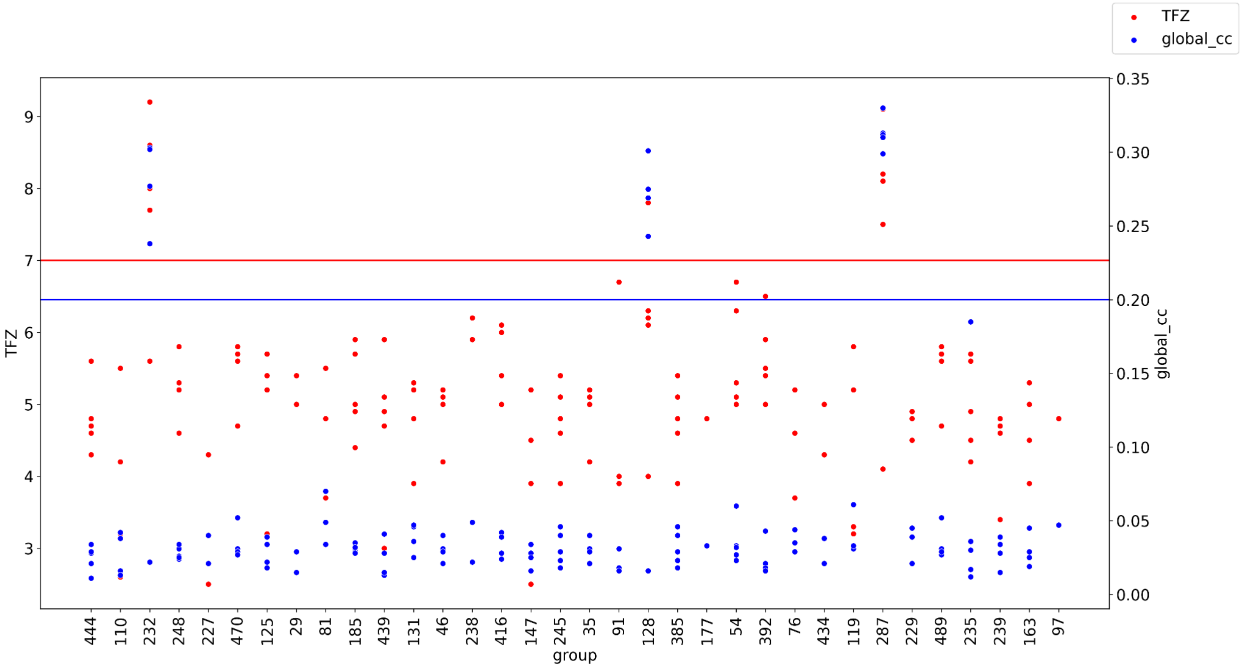


**Supplemental Figure 8. Molecular replacement analysis of all groups for R1117**. LLG (top) and TFZ (bottom) are plotted in red with a horizontal line at 60/7 respectively representing the normal criterion for successful placement. Global Map CC is plotted in blue with a horizontal line at 0.2 representing agreement between the placed model and the solved crystallographic structure.

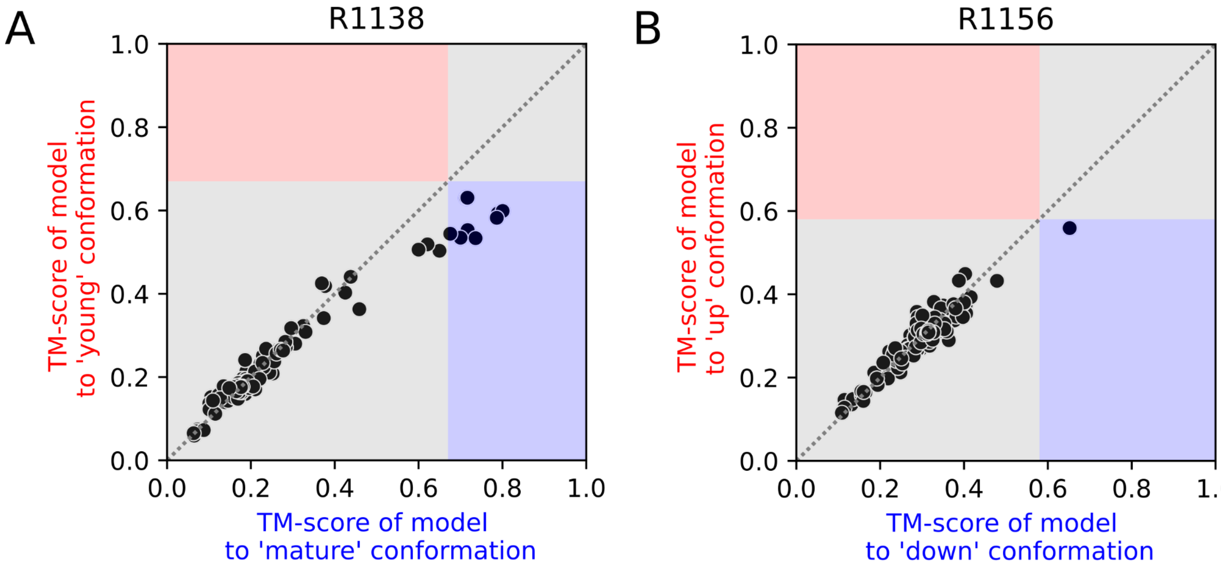


**Supplemental Figure 9. Analysis of how well groups modeled multi-state targets**. For all models submitted, the TM-scores to the two separate conformations are plotted against each other. The two conformations are the mature and young conformations for the 6-helix bundle nanostructure R1138 (A) and 'up' and 'down' conformations for the BtCoV-HKU5 SL5 domain R1156 (B). Gray boxes indicate regions with TM-scores below the TM-score between the two target conformation (0.67 for R1138 and 0.58 for R1156). Red regions indicate models that were close to the young (A) and ‘up’ (B) conformations and blue regions indicate models that were close to the mature (A) and ‘down’ (B) conformations. In both targets, there were models submitted that capture one of the experimentally observed conformations (blue quadrant) but not the other one (red quadrant).
